# High-alpha band synchronization across frontal, parietal and visual cortex mediates behavioral and neuronal effects of visuospatial attention

**DOI:** 10.1101/165563

**Authors:** Muriel Lobier, J. Matias Palva, Satu Palva

**Author notes:** Corresponding authors: Muriel Lobier Helsinki Institute of Life Sciences, Neuroscience Center, University of Helsinki, Finland. Satu Palva Helsinki Institute of Life Sciences, Neuroscience Center, University of Helsinki, Finland.

## Abstract

Visuospatial attention prioritizes processing of attended visual stimuli. It is characterized by lateralized alpha-band (8-14 Hz) amplitude suppression in visual cortex and increased neuronal activity in a network of frontal and parietal areas. It has remained unknown what mechanisms coordinate neuronal processing among frontoparietal network and visual cortices and implement the attention-related modulations of alpha-band amplitudes and behavior. We investigated whether large-scale network synchronization could be such a mechanism. We recorded human cortical activity with magnetoencephalography (MEG) during a visuospatial attention task. We then identified the frequencies and anatomical networks of inter-areal phase synchronization from source localized MEG data. We found that visuospatial attention is associated with robust and sustained long-range synchronization of cortical oscillations exclusively in the high-alpha (10-14 Hz) frequency band. This synchronization connected frontal, parietal and visual regions and was observed concurrently with amplitude suppression of low-alpha (6-9 Hz) band oscillations in visual cortex. Furthermore, stronger high-alpha phase synchronization was associated with decreased reaction times to attended stimuli and larger suppression of alpha-band amplitudes. These results thus show that high-alpha band phase synchronization is functionally significant and could coordinate the neuronal communication underlying the implementation of visuospatial attention.

## Introduction

Attention reconciles the brain’s limited processing capacity with the unlimited flow of sensory input by selecting behaviorally relevant stimuli from irrelevant sensory information. Anticipatory endogenous visuospatial attention improves psychophysical performance in the attended location of the visual field in the absence of eye movement (Posner, 1980) by enhancing neuronal processing of attended stimuli compared to the processing of unattended stimuli in respective sensory cortices (Corbetta and Shulman, 2002; Kastner and Ungerleider, 2000). In electro-(EEG) and magnetoencephalography (MEG) data, anticipatory visuospatial attention suppresses local alpha-band amplitudes more in the visual cortex contralateral to than ipsilateral to the attended hemifield (Capilla et al., 2014; Gould et al., 2011; van Dijk et al., 2008). As alpha suppression is correlated with behavioral performance in both visual and somatosensory tasks with larger suppression being associated with better performance (Iemi et al., 2017; Thut et al., 2006; van Dijk et al., 2008), it is thought to be mechanistically linked to anticipatory attention, possibly through associated changes in neuronal excitability or gain modulations (Iemi et al., 2017; Lange et al., 2013). Local alpha oscillations are thus thought to underlie the inhibition of behaviorally relevant attended information (Jensen and Mazaheri, 2010; Klimesch et al., 2007). Yet, the mechanisms that coordinate the alpha amplitude suppression in sensory cortical areas have remained unknown.

Attentional functions are carried out by the intraparietal sulcus (IPS), superior parietal lobule (SPL), and frontal eye fields (FEF) which together form the dorsal attention network (DAN) (Corbetta and Shulman, 2002; Kastner and Ungerleider, 2000; Petersen and Posner, 2012). The key hubs of this network exhibit mutually correlated blood-oxygenation-level dependent (BOLD) signals (Spadone et al., 2015; Szczepanski et al., 2013). Furthermore, BOLD signal in DAN is biased by visuospatial attention (Szczepanski et al., 2010) and co-varies with attention-related modulations of alpha-band amplitudes in visual cortex (Liu et al., 2016). Finally, the perturbation of neuronal activity in key hubs of DAN with transcranial magnetic stimulation (TMS) modulates the effects of attention on both behavior and alpha-band suppression in visual cortex (Capotosto et al., 2009; Capotosto et al., 2015). DAN is therefore thought to be the brain network responsible for implementing anticipatory visuospatial attention but how neuronal communication is coordinated in DAN and between DAN and the visual system has remained poorly understood.

Neuronal synchronization in the beta (14–30 Hz) and gamma (30–120 Hz) bands has been proposed to coordinate attention-related neuronal processing at sub-second time-scales (Fries, 2015; Miller and Buschman, 2013; Tallon-Baudry, 2012; Womelsdorf and Everling, 2015; Womelsdorf and Fries, 2007). While local field potential (LFP) recordings in non-human primates support this hypothesis (Buschman and Miller, 2007; Gregoriou et al., 2009; Womelsdorf et al., 2007), such strong evidence is lacking for humans. Signal mixing and source leakage hinder reliable estimation of inter-areal synchronization in MEG/EEG sensor-level and source-level analyses respectively (Palva and Palva, 2012; Schoffelen and Gross, 2009). Consequently, only a few prior studies have addressed the role of long-range phase synchronization in attention. Anticipatory visuospatial attention is associated with the lateralization of long-range coherence in the gamma-band between FEF and visual cortex and in the alpha-band between parietal and visual cortex in MEG data (Siegel et al., 2008). Lateralization of alpha-band synchronization between posterior parietal (PPC) and visual cortex during visuospatial attention was also observed in source-reconstructed EEG data (Doesburg et al., 2009). However, as these studies only addressed the differences in synchronization between contra- and ipsilateral hemispheres and were limited to examining small sets of regions-of-interest (ROIs), the complete networks of cortical phase coupling putatively underlying visuospatial attention have remained unidentified and under debate (Palva and Palva, 2007; Palva and Palva, 2011; Sadaghiani and Kleinschmidt, 2016).

We propose here that long-range alpha-band phase synchronization coordinates neuronal processing across relevant cortical areas to support visuospatial attention. We first hypothesized that phase synchronization should be strengthened in task-relevant networks encompassing frontal, parietal and visual cortices during anticipatory visuospatial attention. We further hypothesized that if such synchronization is functionally significant, it should predict attention-related modulations of alpha-amplitude and behavioral performance. To test these hypotheses, we recorded MEG data during a Posner-like cued visuospatial discrimination task. We quantified large-scale network synchronization associated with visuospatial attention using advanced data-analysis techniques and source-localization of the MEG data. To avoid the confounds inherent to frequency- and ROI-limited analyses, we made no *a priori* selection of frequency-bands- or cortical-sources-of-interest. We then identified the most central connections and key cortical areas of significantly strengthened phase synchronized networks using graph theory and investigated their lateralization patterns. Finally, to assess the functional significance of phase synchronization in the coordination of attention, we tested whether its strength co-varied with attention-related modulations of local alpha-band amplitudes and behavioral performance.

## Materials and Methods

### Participants and recordings

Cortical activity was recorded from 14 healthy participants (mean: 26.4 years old, range: 20-33; seven females) with normal or corrected to normal vision using a 306 channel MEG instrument composed of 204 planar gradiometers and 102 magnetometers (Elekta Neuromag, Helsinki, Finland) at 600 Hz sampling rate. After screening of behavioral results, one participant was excluded from further analysis due to very poor performance (13 % detection in the low contrast condition). Maxfilter software (Elekta Neuromag Ltd., Finland) was used to suppress extracranial noise and to co-localize recordings from different sessions and subjects in signal space. Independent component analysis and signal-space projection-based tools were utilized to reject eye-blink, heartbeat and stimulus artifact components from the signal. T1-weighted anatomical MRI scans for cortical surface reconstruction models were obtained for each subject at a resolution of a ≤ 1 x 1 x 1-mm (MP-RAGE) with a 1.5-T MRI scanner (Siemens, Germany). The study was performed according to Declaration of Helsinki and approved by an ethical committee of the Helsinki University Central Hospital. Participants gave written informed consent on participation before the experiment.

### Task

Participants carried out a spatially cued stimulus discrimination task between two possible visual shapes (see Fig. 1) in two different contrast conditions (Low and High). Stimuli were generated using Psychtoolbox-3 (Brainard, 1997). Participants first fixated a red fixation cross (1° of visual angle) centered in a circular patch of dynamic grayscale Perlin noise (diameter =10°). After a jittered duration of 1.25 ± 0.25 s, participants were, on each individual trial, endogenously cued to attend to the right or left hemifield by a change from red to green (calibrated to be equiluminant) of the half of the fixation cross of the hemifield to attend. After a jittered duration of 1.25 ± 0.25 s, one of two geometrical shapes (2°) was presented for 0.1 s in the attended or non-attended lower quadrant. Participants indicated by a thumb lift which shape they had perceived. No response was required if they had not perceived any stimulus. The thumb assigned to each shape was counterbalanced across participants. The fixation cross then changed to blue 2.15 ± 0.25 s after stimulus onset, and participants rated stimulus visibility on a four level scale (not visible, barely visible, partially visible, fully visible) using finger lifts. Participants were instructed to maintain fixation on the fixation cross during the entire trial and eye movements were monitored using EOG. Importantly, participants were asked to respond to both attended and non-attended stimuli. Nine percent of trials were catch trials where no stimuli were presented, 73% were valid trials where the stimulus was presented in the attended hemifield, 18% were invalid trials where the stimulus was presented in the non-attended hemifield, and 9% were catch trials where no stimulus was displayed. Cue type (attend left or right) and trial type (valid, invalid or catch) were pseudorandomized across trials and participants.

**Figure 1.**
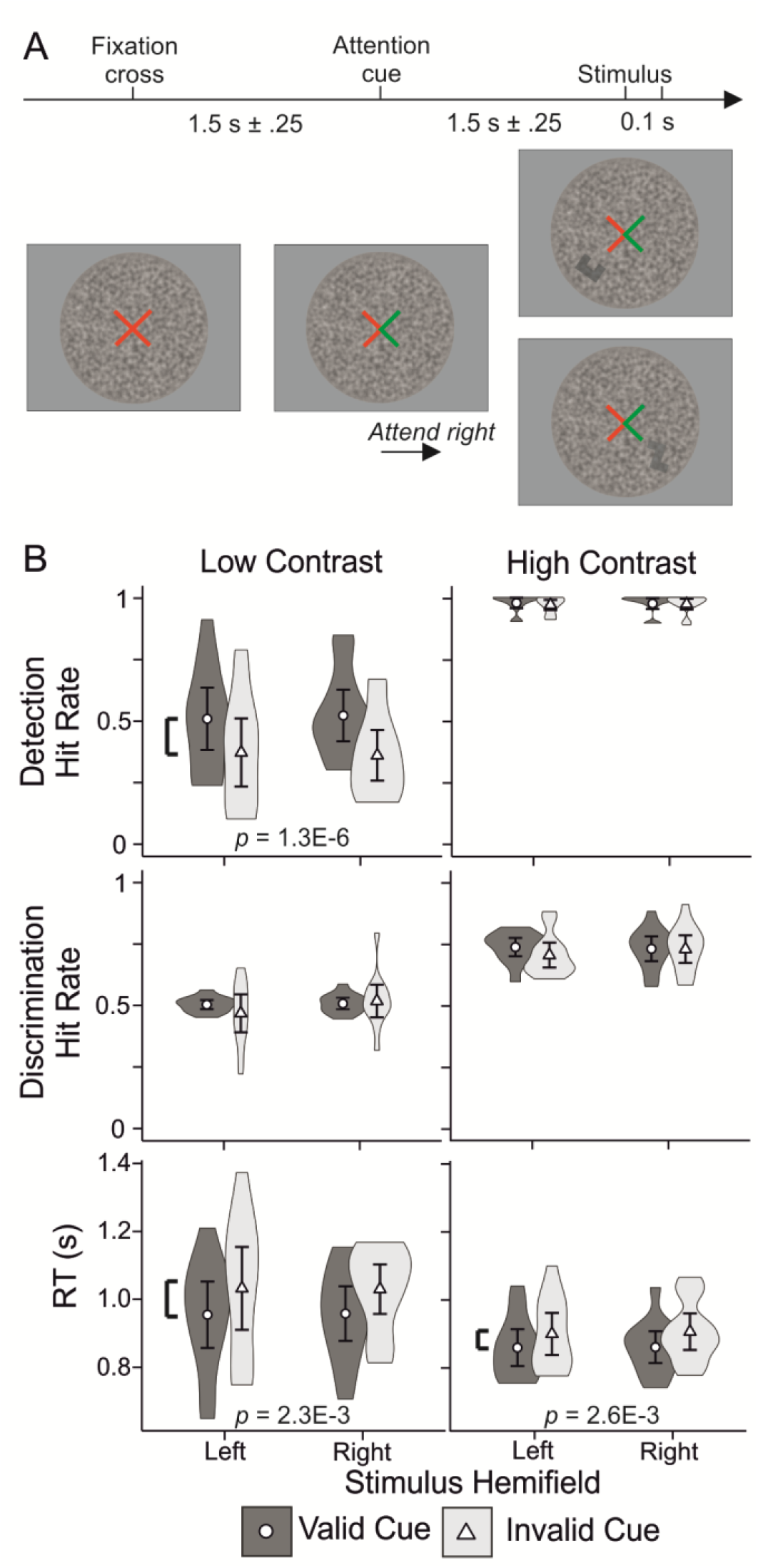
Experimental paradigm and behavioral performance **A** Schematics of the experimental paradigm. Participants were cued to attend to the right or left hemifield by a change from red to green of the relevant half of a central fixation cross. The attention cue remained on screen for the entire duration of the trial. After 1.25 ±.25 s, one of two geometrical shapes was displayed for 0.1 s in the attended or non-attended lower hemifield at either low or high contrast. Participants were asked to discriminate between these two geometrical shapes regardless of the stimulus location. **B** Behavioral performance for detection hit rates (DE-HR, top row), discrimination hit rates (DI-HR, middle row), and reaction times (RT, bottom row) for each contrast (low and high, respectively left and right column) as a function of attend condition (attend-left and attend-right) and cue validity (valid cue and invalid cue). Error bars indicate 95% confidence intervals. P values of significant main effects are provided.

At the beginning of the first recording session, participants carried out two calibration sets of 88 trials each to calibrate individual low and high contrast strengths. The low contrast stimuli were calibrated to a 50% detection rate (participant perceived/detected a shape, irrespective of which one) while the high contrast stimuli were calibrated to a 77% correct discrimination rate (participant perceived/discriminated the correct shape) using the Psychotoolbox implementation of the QUEST algorithm (Watson and Pelli, 1983). Only valid trials were used for calibration, collapsed across attend conditions. Participants then carried out 20 to 25 sets of 88 trials (44 trials for each contrast strength) across four recording sessions. Participants carried out an average of 1053 trials (Range 792-1100) per condition (attend Left – attend Right, collapsed over Low and High contrasts). After rejection of trials contaminated by blinks, eye movements or poor signal to noise ratio, an average of 934 trials per condition (750-1023) remained. In order to avoid effects of sample size bias in our phase synchrony metrics, analyses were run on 750 randomly chosen trials per condition.

### Analysis of Behavioral data

Reaction times (RT), detection hit rates (DE-HR) and discrimination hit rates (DI-HR) were computed for both contrasts. DE-HR was defined as the proportion of detected stimuli, regardless of discrimination accuracy. DI-HR was defined as the proportion of correctly discriminated stimuli. DE-HR and DI-HR were logit-transformed before statistical analysis. To estimate the effect of spatial visual attention on behavioral performance, we ran 2x2 repeated measures ANOVAs with Cue Validity (valid – invalid) and Stimulus Hemifield (Left – Right) as factors on RT, DE-HR and DI-HR data separately for the Low and High contrast conditions.

### Analyses of MEG data

An overview of the workflow including all analysis steps is shown in Inline Supplementary Fig. 1.

#### MEG data preprocessing, filtering, source analysis, and surface parcellations

We used the Maxfilter software (Elekta Neuromag Ltd., Finland) to suppress extra-cranial noise and to co-localize the recordings in signal space. Independent component analysis (ICA, Matlab toolbox Fieldtrip, http://fieldtrip.fcdonders.nl (Oostenveld et al., 2011) was used to extract signal components for the MEG data and to exclude components that were correlated with ocular artefacts identified by EOG or with heart-beat (reference signal was estimated from MEG magnetometers). Source reconstruction and data analysis followed largely previously represented procedures (Palva et al., 2010; Rouhinen et al., 2013) and are here described briefly. Source reconstruction was performed using FreeSurfer software (Fischl, 2012) (http://surfer.nmr.mgh.harvard.edu/) for volumetric segmentation of the MRI data, surface reconstruction, flattening, cortical parcellation, and labeling with the Freesurfer/Destrieux atlas (Destrieux et al., 2010). MNE software (Gramfort et al., 2014) (http://www.nmr.mgh.harvard.edu/martinos/userInfo/data/sofMNE.php) was used to create three-layer boundary element conductivity models and cortically constrained source models for the MEG-MRI co-localization and for the preparation of the forward and inverse operators (Dale et al., 2000). The source models had dipole orientations fixed to pial surface normals and a 5 mm inter-dipole separation throughout the cortex, which yielded models containing 11 000–14 000 source vertices.

We first computed Noise Covariance Matrices (NCMs) using preprocessed broadband filtered (3–40 Hz) MEG time-series from each separate MEG channel. These NCMs were computed from 300 ms time-windows taken in the interval between the visibility response and the start of the following trial. Only time-windows that were not contaminated by eye blink or eye movement artefacts were used for NCM computations. NCMs were then used to compute broadband inverse operators. We then segmented MEG channel time-series into 1.5 second trials spanning from –0.5 s before attention cue onset to 1 s after. We filtered these epoched MEG channel time-series into 32 logarithmically spaced frequencies, *f_min_
* = 3 Hz *… f_max_
* = 120 Hz using Morlet Wavelets with a time-frequency compromise parameter *m* = 5. To reconstruct trial cortical phase-time series, we used individual broadband inverse operators to transform filtered complex single-trial MEG sensor time-series to source-vertex time-series.

These source-vertex time-series were then collapsed into cortical parcel time-series in 400-parcel collections. This parcellation was obtained by iteratively splitting the largest parcels of the Destrieux atlas along their most elongated axis using the same parcel-wise splits for all subjects. Using neuroanatomical labeling as the anatomical “coordinate system” eliminates the need for inter-subject morphing in group-level analyses (which would have compromised individual anatomical accuracy). We performed the source time-series collapsing using sparse weighted fidelity-optimized collapse operators that maximized the reconstruction accuracy in each subject’s individual source space (Korhonen et al., 2014). We computed separately for each frequency-band between-parcel phase synchronization from parcel complex time series *A*(*P*, *n*, *t*)*e*
^
*iφ*(*P*,*n*,*t*)^ where *A(P,n,t)* and *φ(P,n,t)* are the amplitude and phase for parcel P, trial n, and time sample t.

##### Analysis of inter-parcel phase synchrony

To identify cortex-wide phase synchrony networks, we first computed individual parcel-parcel interaction matrices for each condition, frequency and 0.2 s time-window with a 0.1 s overlap from –0.5 s to 1 s with both the weighed Phase Lag Index (wPLI) (Vinck et al., 2011) and the Phase Locking Value (PLV) (Lachaux et al., 1999). We chose wPLI because of its lack of sensitivity to zero phase-lag interactions. These zero-phase lag artificial, non-true interactions are a consequence of the remaining signal mixing between parcel-time series after source reconstruction (Palva and Palva, 2012). WPLI synchrony matrices should therefore be free of artificial parcel-parcel interactions at the cost of missing any true zero-phase lag interaction. To ensure that our pattern of results was not biased by this limitation, we thus also computed PLV which is equally sensitive to synchrony at all phase-lags. We collapsed the 400 parcel data to a coarser parcellation of 200 parcels before computing interaction metrics.

To compute inter-parcel phase synchrony metrics, we used the complex parcel time series for each frequency and condition. For each time-window, frequency and condition, we computed *PLV* between parcels P_1_ and P_2_ for N trials and T time samples per trial as

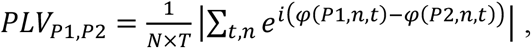

where *φ*(*Pk*, *t*, *n*) is the phase of parcel *Pk* in trial *n* and time sample *t*. We computed wPLI as

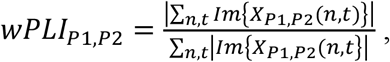

where *X_P1,P2_
* is the cross-spectrum of the complex time-series of parcels P_1_ and P_2_ computed as *X*
_
*P*1,*P*2_ = *A*(*P*1, *n*, *t*)*A*(*P*2, *n*, *t*)*e*
^
*i*(*φ*(*P*1,*n*,*t*)−*φ*(*P*2,*n*,*t*))^. The synchrony data therefore comprised two 200 x 200 parcel-parcel interaction matrices (one for each metric) for each combination of 13 subjects, 2 conditions, 32 frequency bands and 14 time-windows. Phase-synchrony metrics were baseline corrected by subtracting the mean baseline phase synchrony before statistical testing (computed by averaging phase synchrony values across time-windows between –0.5 and –0.1 s for each frequency and condition).

Group statistics were performed separately for each frequency and time-window to identify significant parcel-parcel interactions. Screening of behavioral results revealed that two participants did not exhibit a consistent behavioral effect of visuospatial attention (decrease in RTs and increase in HR). Group contrasts analyses were thus carried out on the remaining eleven participants. Significance thresholds were estimated using a Wilcoxon Signed Rank test with *α* = 0.05. To reduce the false discovery rate in the performed multiple statistical comparisons, for each contrast we pooled significant observations over all parcel-parcel interactions and then discarded as many least-significant comparisons as were predicted to be false discoveries by the alpha-level used in the corresponding test (Palva et al., 2010; Palva et al., 2011). For each metric, the synchrony data was therefore reduced to two group-level adjacency matrices for each combination of condition, frequency and time-window: one for the positive tail (increase in phase synchrony) and one for the negative tail (decrease in phase synchrony). In these adjacency matrices, the value of non-significant interactions was zero and the value of significant interactions was the average baseline-corrected interaction strength across participants. To further control for false discoveries, if the fraction of significant interactions within a given adjacency matrix was inferior to 0.002, all interactions were set to zero.

##### Removal of low-fidelity – high infidelity parcels and connections

A major limitation of connectivity analysis using MEG data is signal linear mixing between neighboring parcels that remains after source reconstruction (Palva and Palva, 2012). This signal mixing is more or less prevalent depending on source anatomical location and individual anatomy. *Fidelity* (Korhonen et al., 2014), uses simulated data to capture how well the original time-series of a parcel is represented by the reconstructed time-series of that same source parcel. It is defined as the phase correlation (real part of the PLV) between the original simulated time-series of a parcel and its forward-inverse modeled (reconstructed) time-series. In parcels with low *fidelity*, the phase of the forward-inverse modeled parcel time-series is poorly correlated with the phase of original simulated parcel time-series, suggesting low reconstruction accuracy. *Infidelity*, on the other hand, captures how much the reconstructed parcel time-series is contaminated by the original time-series of neighboring parcels. In parcels with high infidelity, the phase of the forward-inverse modeled parcel time-series is moderately correlated with the phase of the original simulated time-series of neighboring parcels. To avoid spurious or misplaced interactions, we removed from the adjacency matrices parcels whose *fidelity* is in the bottom 5 percent or whose *infidelity* is in the top 5 percent. The eighteen parcels thus selected were mostly deep and/or inferior sources, which generate the least detectable signals and thus are most likely to incorrectly reflect signals generated elsewhere. In addition to removing very unreliable parcels (and thus all their possible parcel-parcel interactions), we removed parcel-parcel interactions that connected two moderately low *fidelity* parcels. In total, we removed 24% of interactions as likely to be contaminated by source-level linear mixing. To visualize the cortical distribution of edge removal, we mapped the proportion of removed interactions for each parcel on the surface of a 3D inflated brain in Inline Supplementary Fig. 2.

##### Network visualizations

We used graph theory (Bullmore and Sporns, 2009) to characterize group-level adjacency matrices. Each thresholded adjacency matrix defined a graph made up of nodes and edges, where nodes are parcels that have at least one significant interaction and edges are the significant interactions between nodes. Connection density (*κ)* indexes the proportion of edges (significant interactions) from all possible interactions. Graph strength (*GS*) is the summed strength of all edges. Node strength (*NS*) is the summed strength of all edges connected to a given node. To identify nodes that play a central role within each graph, we computed node Eigen Vector Centrality (*EVC*) (Bonacich, 2007). *EVC* takes into account both the number and the ‘quality’ of connections to a given node: an edge connected to a node with high *EVC* will be weighed more heavily than an edge connected to a node with low *EVC*. As a result, a node with few edges to high-*EVC* nodes may have a higher *EVC* than a node with more edges to low *EVC* nodes. To identify edges that play a central role within each graph, we defined Edge *EVC* as the sum of the *EVC* of the edge’s two nodes.

To investigate the time and frequency patterns of phase synchrony modulations associated with visuospatial attention, we first computed a compound time-frequency plot for inter-parcel phase synchrony for each attend condition and phase synchrony metric. We plotted the connection density *κ* for both the positive tail graphs (*κ+*, increase in inter-parcel synchrony compared to baseline) and the negative tail graphs (*κ-*, decrease in inter-parcel synchrony compared to baseline) of each TF bin.

Graph visualization was carried out for frequency-bands and time-windows showing significant increases in phase synchrony. For each selected frequency band, we first constructed a single graph for each condition and time-window by summing the adjacency matrices of each filtering frequency included in the frequency band. We then picked the 500 most influential edges from these graphs as defined by Edge *EVC*. To take into account the presence of spurious edges (‘false’ interactions created by the concurrent presence of a true interaction and linear mixing, (Palva and Palva, 2012), we then applied a novel edge-bundling approach.

The reconstructed time-series of parcels close to the truly connected parcels are contaminated by signals that are truly phase synchronized, therefore, these parcels will also appear to be phase synchronized. Spurious interactions are thus present between parcels close to true interactions. To minimize the impact of these spurious interactions, we used an edge bundling method that clusters together all edges that are likely to be manifestations of the same true interaction. We first estimated the ‘edge proximity’ of each edge pair as a function of the linear mixing between the nodes/parcels of each edge. If there is high linear mixing across the parcels that define two separate edges, it is more likely that these two edges reflect the same true interaction than if the linear mixing is low.

We used infidelity (Korhonen et al., 2014) as a measure of the amount of linear mixing between two parcels. Parcel pairs with high infidelity are more contaminated with each other’s signal than parcel pairs with low infidelity. For a given pair of edges X (linking parcels X_1_ and X_2_) and Y (linking parcels Y_1_ and Y_2_), we then define edge proximity as *max*(*infid*(*X*
_1_, *Y*
_1_) × *infid*(*X*
_2_, *Y*
_2_), *infid*(*X*
_1_, *Y*
_2_) × *infid*(*X*
_2_, *Y*
_1_)). We then applied hierarchical clustering to ‘edge proximity’, in order to cluster or ‘bundle’ together the edges most likely to reflect the same interaction. The size of the bundle reflected the spatial uncertainty of the exact location of the true interaction it represents. To draw the graphs, we used lines to indicate edges, *i.e*., significant interactions, and circles to denote the nodes, *i.e*., cortical parcels. Graphs were overlaid on flattened maps of the complete cortical surfaces of left and right hemispheres described previously. Color in flattened brains identifies the 7 brain systems of the Yeo parcellations (Yeo et al., 2011) and the thin white lines the 148 parcels of the Destrieux parcellation (Destrieux et al., 2010). Node size was proportional to node *EVC*. Node names were displayed for nodes belonging to relevant brain systems (visual, dorsal attention network DAN, ventral attention network VAN, frontoparietal) and to the 20% most influential nodes (as defined by node *EVC*). We excluded bundles of less than four edges as more likely to be false positives. Finally, in each edge bundle thought to reflect the same true interaction, we only drew the 80 % edges with highest proximity.

##### Analysis of local oscillation amplitudes and stimulus phase-locking

To investigate whether the increase in high-alpha phase synchrony was associated with a specific response in local oscillation amplitudes, we computed the amplitude time-frequency response across all cortical parcels and summarized the results in a time-frequency plot. We computed within-parcel amplitude envelopes (equivalent to the square root of oscillatory power) by averaging within-parcel amplitudes of the Morlet-filtered time-series across *N* trials and *T* samples for each condition and time-window: 
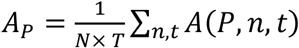
In addition we computed within-parcel stimulus phase-locking (SL) – phase-locking of ongoing oscillations to stimulus onset – by taking the norm of the averaged within-parcel phase values of the Morlet-filtered time-series across *N* trials and *T* samples for each condition and time-window: 
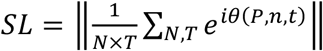
. Trial-averaged oscillations were thus decimated into mean amplitudes and SL within thirty 100 ms long time-windows with a 50 ms overlap from –500 ms to 1000 ms, 0 being attention cue onset. We collapsed the 400 parcel data to a coarser parcellation of 200 parcels. The data therefore comprised a volume of 13 subjects, 200 parcels, 2 attend conditions, 32 frequency bands and 30 time-windows. Before statistical testing, amplitude data were baseline corrected by computing the logarithm of the baseline corrected amplitude: 
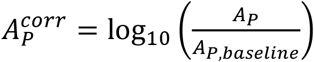
 and SL data were baseline corrected by subtracting the mean baseline SL. We computed baseline amplitudes/SL by averaging amplitudes/SL across time-windows between −500 and −100 ms for each frequency and attend condition. Similar to the phase synchrony analysis, the two participants showing no behavioral effects of visual attention were excluded from group contrasts.

Group statistics were performed separately for each frequency and time-window. Significant differences in parcel oscillation amplitudes (intra-parcel amplitude data) were estimated using a Wilcoxon Signed Rank test with *α* = 0.05. False discovery rate was reduced as described previously but with pooling of significant observations over all cortical parcels. In addition, for a given parcel, we only considered as significant time-frequency bins that were statistically significant in three consecutive overlapping time-windows (i.e., 200 ms).

We first visualized amplitude modulations across frequencies using amplitude time-frequency (TF) plots for each condition (attend left-attend right). For each TF bin, we computed the fraction of cortical parcels that showed a significant amplitude increase (*i.e*., *P_TF_
*+, the fraction of significant positive parcels for each TF bin) or decrease (*i.e*., *P_TF_
*–, the fraction of significant negative parcels for each TF bin) compared to baseline.

We next explored the localization of amplitude modulations induced by visual attention in separate TF windows of interest (6-9 Hz and 400-600ms, 600-800ms and 800-1000 ms). For each time-frequency window, we computed, for each parcel, the fraction of TF bins that had shown a significant amplitude decrease compared to baseline in the previous statistical analysis (*i.e*., *P_P_–*, the fraction of significant negative TF windows for each parcel). We then displayed P_P_- on an inflated 3D brain, therefore highlighting cortical parcels where the decrease in oscillation amplitude was most consistent for each TF window of interest. In addition, we computed for both low (6-9 Hz) and high (11-14 Hz) alpha-bands the average significant amplitude modulation for each one of the previously specified time-windows. These average amplitude modulations were then displayed on inflated 3D brains, highlighting those areas with the strongest modulations of oscillation amplitudes.

##### Correlation between low-alpha amplitude response and low-alpha inter-areal phase synchronization response

We investigated whether modulations in low-alpha amplitudes could explain similar modulations of low-alpha wPLI, i.e., if a stronger decrease in local low-alpha amplitudes could be associated with a stronger decrease in low alpha wPLI because of the resulting signal to noise ratio (SNR) decrease. First, we estimated, for each participant, the correlation between low-alpha amplitude and wPLI network strength across parcels in three separate time-windows of interest, 400–600 ms, 600–800 ms, and 800–1000 ms. For each parcel, wPLI strength was computed as the sum of the wPLI strength of all connections to this parcel. Furthermore, wPLI strength was computed before any statistical thresholding, therefore the only edges (connections) removed were those identified as unreliable as presented in section 2.4.3. Correlations were thus computed across the 182 parcels having at least one possible edge/connection. We then used the resulting three samples (one for each time-window) of thirteen correlation coefficients (see Inline Supplementary Fig. 6) to test for the presence of a non-zero correlation. We first Fischer-transformed the correlation coefficients and then applied, in each time-window, a one-sample t-test against 0. We used an *α*-level of 0.05 to determine significance and applied Holms-Bonferroni correction to account for multiple comparisons.

### Lateralization patterns of high-alpha phase synchrony and low-alpha amplitudes

We investigated the lateralization patterns of high-alpha phase synchronized networks as well as of low-alpha visual cortex oscillation amplitudes. We therefore investigated whether high-alpha interareal phase synchrony was lateralized similarly to the expected lateralization of low-alpha local amplitudes. For each participant we first computed ipsi- and contralateral intra-hemispheric graph strength of high-alpha band synchronization for each time-frequency bin and attend condition. We then averaged, for each participant, low-alpha baseline-corrected amplitudes across all cortical parcels belonging to ipsi- or contralateral visual cortex (parcels included in visual cortex are illustrated in Inline Supplementary Fig. 3) for each time-frequency bin and attend condition. We then investigated the effect of visuospatial attention on the hemispheric balance (contralateral versus ipsilateral) of alpha amplitudes in visual cortex. We first computed the average low-alpha band amplitude response in visual cortex by averaging low-alpha band amplitudes across visual cortex parcels (as defined by a morphing of the Destrieux and Yeo parcellations) for each TF bin and hemisphere. We thus obtained, for each attend condition, two average amplitude time-series

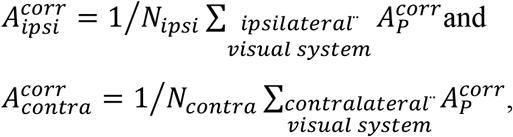

where *N_ipsi_
* and *N_contra_
* are the number of parcels mapped to the ipsilateral and contralateral visual system. We then investigated statistically significant differences between ipsi- and contralateral low-alpha amplitudes during visuospatial attention. To reduce the number of statistical comparisons, we averaged the data in non-overlapping 200 ms time bins spanning 0 to 1 s post cue onset. For each attend condition and time bin, we then computed Wilcoxon Signed Rank tests. We used an α-level of 0.05 to determine significance and applied Holms-Bonferroni correction to account for the ten comparisons we carried out.

Finally, we explored the lateralization of the high-alpha phase synchronization networks within the visual system and between the visual and DAN systems. For each attend condition, we counted the number of significant connections within ipsi- and contralateral visual cortex as well as between ipsi- and contralateral visual cortex and bilateral DAN. We computed a lateralization index (LI) as 
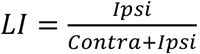
, where values above/below 0.5 indicate more/less connectivity in the ipsilateral compared to the contralateral hemifield. We obtained the null-hypothesis distributions for this LI by random shuffling of the networks. For each attend-condition and time-window, we randomly shuffled the significant edges in the network across all possible edges and then computed the LI values for Visual-Visual and Visual-DAN connectivity. We carried out 1000 shuffles for each condition, and from these distributions determined the 95% confidence limits on the null-hypothesis. We then considered connectivity to be more strongly lateralized than would be expected by chance when LI values did not fall into these 95% confidence limits.

### Relationship between low-alpha amplitude lateralization and high-alpha inter-areal phase-synchronization

To explore the putative links between attention related modulations of high-alpha-band inter-real phase synchrony and low-alpha band oscillation amplitudes, we tested whether graph strength in the high-alpha band and amplitude modulations in the low-alpha band co-varied across all thirteen participants.

For each attention condition (Attend Left and Attend Right) and time-window (400-600 ms, 600-800 ms, 800-1000 ms), we computed individual amplitude lateralization indices (*ALI*) for parcels mapped to the visual system according to our morphing of the Destrieux and Yeo parcellations as well as phase synchronization graph strength (*GS*). *ALI* was computed as the difference between the average low-alpha band baseline-corrected response in contralateral visual system parcels and ipsilateral visual system parcels:

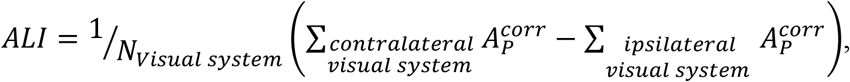

where *N_visual system_
* is the number of parcels mapped to the visual system. More negative *ALI* values therefore indicate stronger alpha suppression in contralateral compared to ipsilateral visual cortex. To estimate individual *GS* values, we first computed individual weighed graphs by multiplying individual baseline corrected wPLI interaction matrices by a binary mask based on group graphs. For each condition, edges that were not significant at the group level were set to zero. We then computed individual *GS* for each attend condition and time-window from these individual weighed graphs as *GS* = ∑_
*all edges*
_ *wPLI*. In each time window, we then collapsed the data from both attend conditions. To account for the resulting non-independence between data points, we used R (R Core Team, 2016) and lme4 (Bates et al., 2015) to perform linear mixed effects analyses. For each time-window, we built a full model and a null model. In the full model, we entered GS as a fixed effect. For random effects, we had intercepts for participants. In the null model, we only included participants as random effects. *P*-values were obtained by likelihood ratio tests of the full model (which included the fixed effect of interest, GS) against the null model (which did not include the fixed effect of interest). *P*-values were subsequently corrected for multiple comparisons using Holms-Bonferronni correction. The amount of variance in ALI explained by GS in the full model was estimated using the marginal R-squared computed using the MuMin R package.

### Correlation between individual graph strength and behavioral performance

We investigated whether individual network strength co-varied with the effect of visuospatial attention on individual behavioral performance across all thirteen participants. To quantify the behavioral effect of visuospatial attention, we computed a normalized RT benefit (*RT_benefit_
*) for each participant, contrast strength and attention condition as

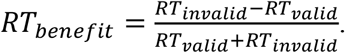

A positive *RT_benefit_
* indicates faster responses for attended than unattended stimuli. We then computed individual *GS* values separately for each contrast and attend condition by averaging *high-α* GS across all three time windows (400–600, 600–800, 800–1000 ms). To test whether stronger average *GS* was associated with a larger behavioral benefit of visuospatial attention (i.e., for a positive correlation), we collapsed data across participants separately for each contrast strength and attend condition. We then used the MATLAB Robust Correlation Toolbox (Pernet et al., 2013) to compute both Pearson (*r*) and Spearman (*ρ*) correlation coefficients as well as bootstrapped confidence intervals to evaluate whether average *GS* was correlated with *RT_benefit_
*. Both *p*-values and confidence intervals were Bonferonni corrected for multiple comparisons.

## Results

### Visual attention improves behavioral performance

We measured MEG data from 14 participants performing a cued visuospatial attention task. Participants were first cued to attend to the left or right visual hemifield after which one of two target shapes was displayed in either hemifield at high (75% discrimination) or low (50% detection) contrast (Fig.1A). They discriminated and reported shape identity regardless of its location. In the Low Contrast condition, DI-HR was at chance level (~0.5) while in the high contrast condition, DE-HR was at ceiling (~1), and therefore statistical analyses were not performed on these conditions. The mean false-alarm rate was 0.035 (Range = .006-0.1). Results are presented in Fig. 1B.

In the Low Contrast condition, visuospatial attention had similar effects on RTs and DE-HRs. Participant responded faster and detected stimuli better for attended (valid trials) than unattended (invalid trials) stimuli (*N* = 13, RTs: *F*(1,12) = 14.8, *p* = 2.3E-3, *η^2^
* = .06, DE-HR: *F*(1,12) = 78, *p* = 1.3E-6, *η^2^
* = .15). Stimulus hemifield did not affect performance and no interaction of cue validity with stimulus hemifield was observed (*N* = 13, *F* < 1, *p* > .7 in all cases). In the High Contrast condition, visual attention had different effects on RTs and DI-HRs. Participants responded faster for attended compared to unattended stimuli (*N* = 13, *F*(1,12) = 14.3, *p* = 2.6E-3, *η^2^
* = .06), but did not discriminate attended better than unattended stimuli (*N* = 13, *F*(1,12) = 1, *p* = 0.34). Similar to the low contrast condition, stimulus hemifield did not affect performance and no interaction of cue validity with stimulus hemifield was observed (*N* = 13, *F* < 1, *p* > .2 in all cases). For high contrast stimuli, anticipatory visuospatial attention thus improved reaction times but not discriminability. These results validate that participants were covertly attending to the cued hemifield.

### Sustained visuospatial attention is associated with a sustained increase in inter-areal phase synchrony in the high-α frequency band

Our central aim was to identify the most prominent modulations of inter-areal synchronization by visuospatial attention. We thus estimated the strength of phase synchronization between all cortical parcel pairs using for frequencies between 3–120 Hz and identified connections with statistically significant modulations of synchronization compared to baseline (Wilcoxon Signed Rank test, *α* = 0.05, FDR-reduced). We then used connection density *K* (Rubinov and Sporns, 2010), *i.e*., the proportion of significant connections out of all possible connections, to quantify the extent of synchronization. We summarized these data into time-frequency plots (TF-plots).

TF-plots showed similar spectro-temporal patterns of inter-areal synchronization for attend-left and attend-right conditions. Inter-areal phase synchrony, as estimated with wPLI (Fig. 2A) or PLV (Inline Supplementary Fig. 4), was strengthened transiently between 0 and 0.5 s after cue onset in the theta (3–7 Hz) frequency band. This transient theta-band synchronization was followed by a sustained strengthening in high-alpha (11–14 Hz) band phase-synchronization starting approximately 400 ms after cue onset. High-alpha synchronization was observed with both wPLI and PLV, and therefore should reflect strengthening of neuronal phase synchrony that is neither attributable to signal mixing (wPLI) nor to systematic changes in phase lags without a change in coupling strength (PLV). Importantly, sustained inter-areal synchronization was not present in any other frequency band. Suppression of synchronization in the low-alpha (6–9 Hz) frequency band was however present, albeit only for wPLI (Fig. 2A and Inline Supplementary Fig. 4).

**Figure 2.**
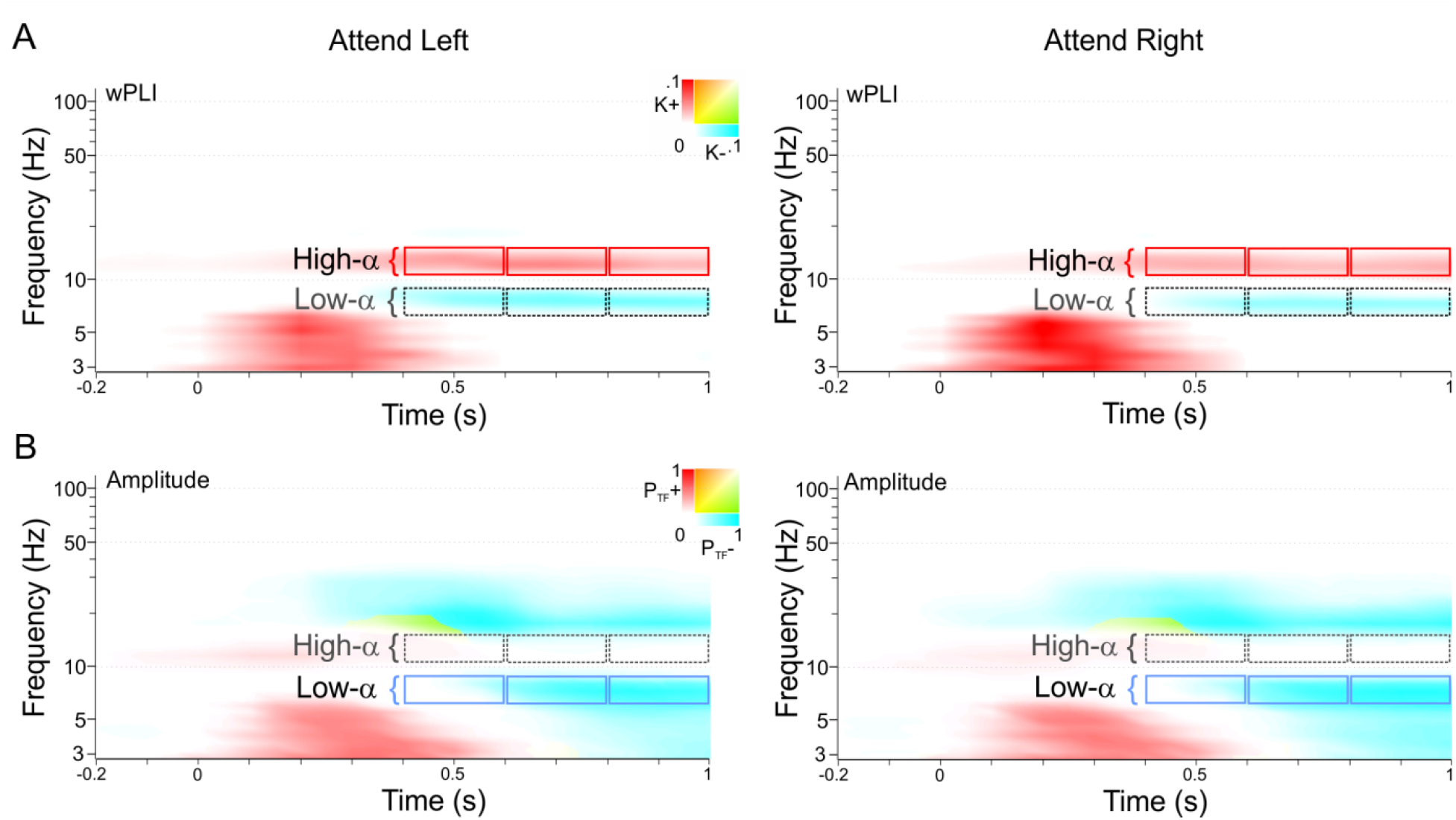
*Visuospatial attention modulates high-alpha band inter-areal phase and low-alpha local oscillation amplitudes*. **A** Time-frequency plots of significant modulations of inter-areal phase synchrony between cortical parcels induced by visuospatial attention as estimated with wPLI. Color indicates connection density (K). K+ represents the connection density for increases in phase synchrony compared to baseline while K-represents its decreases. **B** Time-frequency plots of the proportion of cortical parcels with significant enhancement (P+) or suppression (P-) of local oscillation amplitudes for the attend-left and attend-right conditions.

### High-alpha band synchrony networks connect frontal, parietal and visual cortices

High-alpha band phase synchronization should connect task-relevant cortical regions if it were to be functionally significant in visuospatial attention. We thus mapped the anatomical structure of high-alpha (11–14 Hz) phase synchronized networks in three representative 200 ms time-windows (TWs) (400–600 ms, 600–800 ms and 800–1000 ms from cue onset, red outlines in Fig. 2A). To facilitate interpretation, we only considered the 500 most central edges (according to eigenvector centrality (EVC)) within and between relevant cortical systems (visual, dorsal attention network (DAN), ventral attention network (VAN) and frontoparietal network (FPN) (Fig. 3). The distributions of node and edge EVCs are provided in Inline Supplementary Fig. **5.**

**Figure 3.**
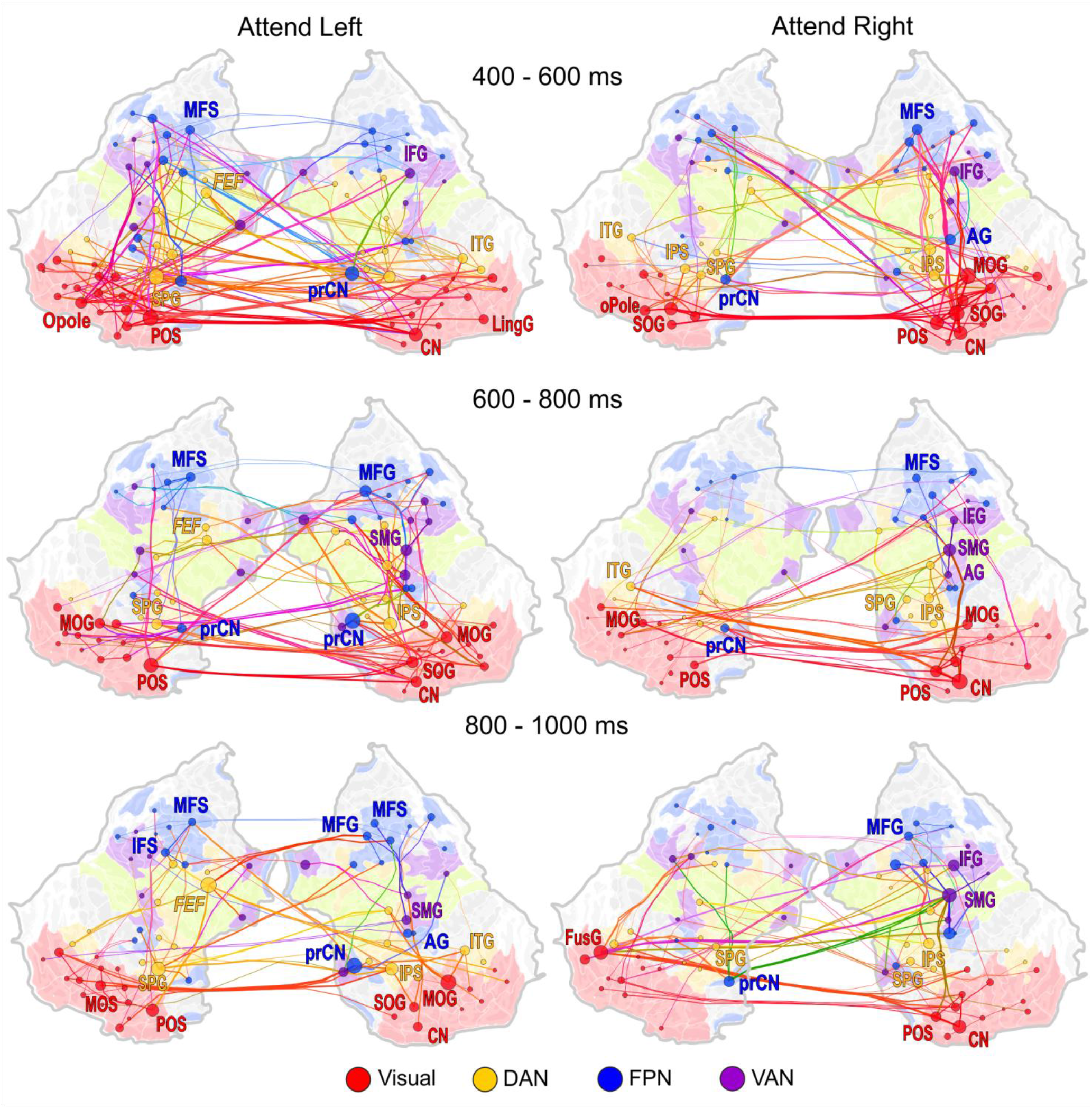
*Networks of high alpha-band inter-areal phase synchronization*. Significant connections of high-alpha band synchronization for three different time-windows (400-600 ms, 600-800 ms and 800-1000 ms) estimated with wPLI and displayed on an inflated and flattened cortical surfaces for attend-left and attend-right conditions (left and right column respectively). Figures display edges that (1) are amongst the 500 highest Edge Eigenvector Centrality (ECV) edges and (2) belong to the visual, DAN, FPN or VAN systems. Node size represents node ECV while the color of nodes, edges, and cortical surfaces represent different functional systems as indicated below. DAN = dorsal attention network, VAN = ventral attention network, FPN: frontoparietal. Node abbreviations are as follows: cuneus (CN), posterior occipital sulcus (POS), superior/middle occipital gyrus (S/MOG), middle occipital sulcus (MOS), fusiform gyrus (FusG), superior parietal gyrus (SPG), intraparietal sulcus (IPS), inferior temporal gyrus (ITG), frontal eye fields (FEF, located in the superior precentral sulcus), precuneus (prCN), angular gyrus (AG), middle frontal sulcus/gyrus (MFS/MFG), inferior frontal sulcus (IFS), supramarginal gyrus (SMG), inferior frontal gyrus (IFG)

High-alpha band phase synchronization was observed both within and between visual and frontal and parietal cortices. During the first TW, the most central hubs for high-alpha band phase-synchronization were observed in cuneus (CN), occipital pole (Opole), and lingual gyrus (LinG) corresponding to V1 as well as in parieto-occipital cortex (POS) and superior occipital gyrus (SOG) of the lateral occipital cortex (LOC). Synchronization connected hubs in the visual system both between hemispheres and within hemispheres. Central network hubs were also found in right intraparietal sulcus (IPS), right inferotemporal gyrus (ITG), left superior parietal gyrus (SPG), and left frontal eye fields (FEF) of DAN, contralateral precunei (prCN), middle frontal sulcus (MFS), and bilateral right angular gyrus (AG) of FPN as well as in the inferior frontal gyrus (IFG) of VAN. Importantly, high-alpha band phase synchronization amongst these key hubs connected the visual system, DAN, and FPN. MFG and FEF of the prefrontal cortex (PFC) were also respectively coupled with CN / POS and ITG of the visual system.

In the second and third TWs, the most central hubs were similar to those of the first time-window. In the visual system, they were observed in CN, POS, SOG and MOG bilaterally for the attend-left condition and ipsilaterally for the attend-right condition. For both time-windows and attend-conditions, hubs were present in DAN (right IPS, ipsilateral SPG, and left FEF), FPN (MFS and prCN), and VAN (left IFG). Additional nodes included right ITG for the attend-left condition and left ITG for the attend-right condition. However, the connectivity patterns between these hubs and cortical systems differed from the first time-window. In the visual system, within-hemisphere edges were present bilaterally in the second TW but only ipsilaterally in the third TW. While FEF was connected to ipsi- and contralateral visual systems by influential edges in the last time-window for the attend-left condition, overall edges between DAN hubs and the visual system were fewer and less central. Similarly, connectivity within attention systems, *i.e*., between FPN hubs (such as MFS and MFG) and DAN/VAN hubs, was more frequent and more central than connectivity between MFS/MFG and the visual system.

To ensure that the increase in phase synchronization did not result from widespread and sustained phase locking of local oscillations to the stimuli (stimulus locking, SL), such as “alpha-ringing” (Makeig et al., 2002), we computed the TF-plot of SL values (Inline Supplementary Fig. 4.B). As expected in an event-related paradigm, SL displayed a strong but transient increase between 0 to 600 ms after the cue-onset in low-frequencies (3-12 Hz). This increase in SL, however, occurred earlier and in lower frequencies than high-alpha phase synchronization, and therefore cannot explain the observations of large-scale high-alpha phase synchrony.

To provide an overall view of these cortical networks, we also visualized the top 400 edges across all systems (Visual, DAN, VAN, FPN, somatomotor (SM), default mode network (DMN), and limbic) in Inline Supplementary Fig. 6. In addition to phase synchrony within and between attention networks and the visual system, FPN and DAN were connected to the SM system, especially during the later time-windows. Such connectivity within SM may reflect anticipation for the response to be made after the stimulus presentation.

### Anticipatory visuospatial attention is associated with low-alpha oscillation amplitude suppression

We then investigated whether visuospatial attention was associated with the commonly observed alpha-band amplitude suppression. We computed the average oscillation amplitudes separately for each time-frequency (TF) bin and cortical parcel and estimated the fraction, *P*, of cortical parcels with statistically significant (Wilcoxon Signed Rank test, α = 0.05, FDR-reduced) modulations of oscillation amplitudes compared to pre-cue baseline (Fig. 2B). Modulations of cortical oscillation amplitudes were similar for attend-left and attend-right conditions. An early (100–500 ms post attention cue) increase in theta band amplitudes was followed by a decrease in oscillation amplitudes 500 ms after attention cue onset. This amplitude suppression was strongest in the low-alpha (6–9 Hz) band but was also present in the beta (18–30 Hz) and theta-bands. For each time-window of interest previously defined, we computed, within the low-alpha band, the proportion of TF bins for each parcel with a significant decrease in amplitudes P-(Fig. 4). Low-alpha band suppression was more extensive in the contra-than ipsilateral visual cortex. Amplitudes were suppressed not only in task-relevant sensory cortex but also in frontal areas such as SFG and MFG of LPFC as well as in IPS and SPL of PPC. Suppression was most pronounced in contralateral middle occipital cortex (middle occipital gyrus and sulcus, MOG and MOS) for both attend-right and attend-left conditions. The TF plot in Fig. 2B shows also that the widespread amplitude suppression in the low-alpha band did not extend to the high-alpha (11–14 Hz) band where large-scale phase synchronization was found. The absence of concurrent widespread changes in high-alpha band amplitudes during high-alpha band phase synchronization precludes the possibility that changes in signal-to-noise (SNR) could underlie the observed changes in the strength of synchronization.

**Figure 4.**
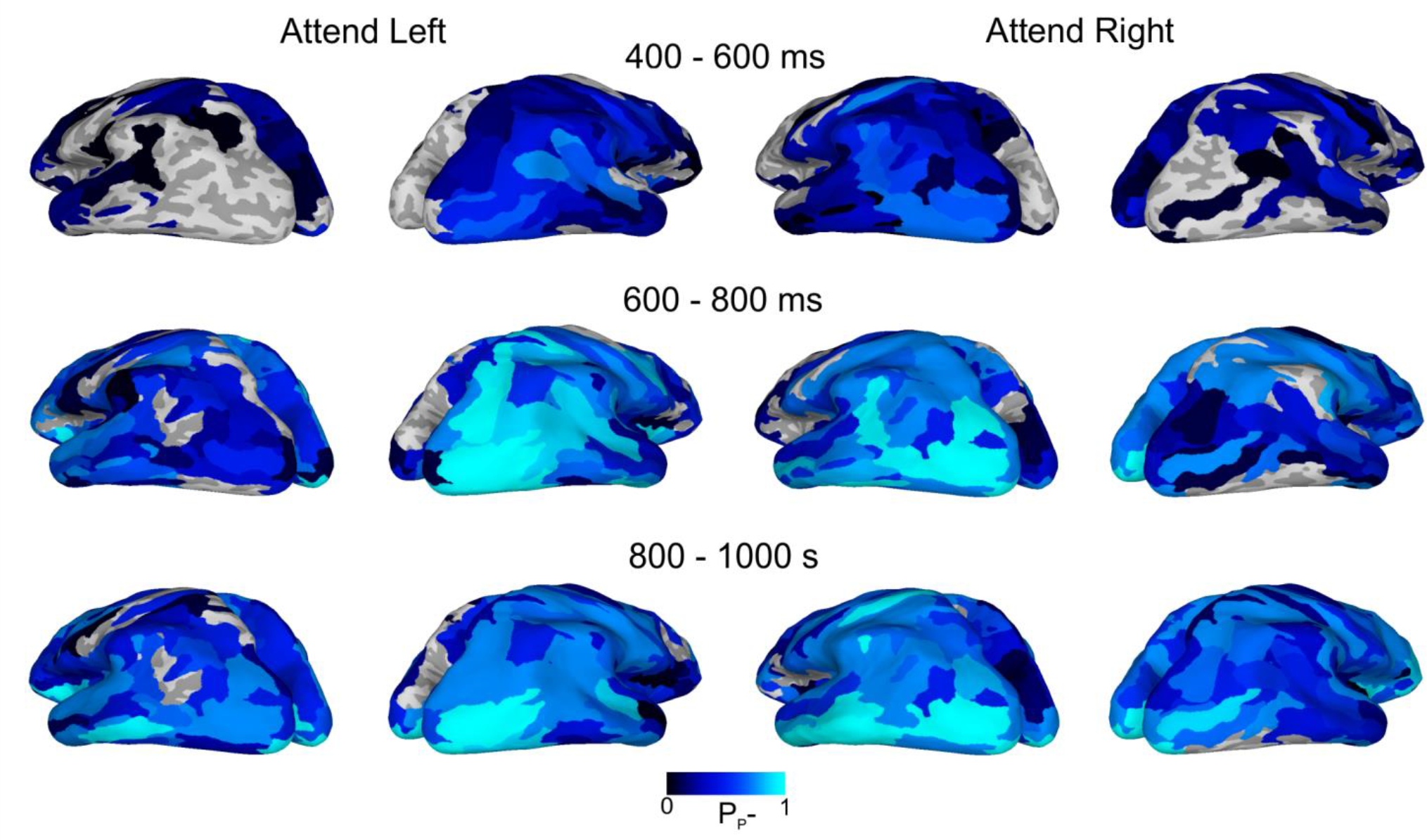
*Cortical localization of the low-alpha band amplitude suppression*. Parcel-wise fraction of significantly negatively modulated TF bins (P-) in the 6-9 Hz frequency range for three different time-windows, displayed on an inflated three dimensional cortical surface.

To confirm this observation, we computed, for each time-window of interest, the average low-alpha and high-alpha significant amplitude modulations for each parcel (Inline Supplementary Fig. 7). As suggested by Fig. 4, low alpha amplitude suppression spread beyond the visual cortex and was stronger in contra-than ipsilateral cortex. In contrast, high alpha amplitudes modulations were restricted to visual cortex and were both suppressed and enhanced relative to baseline levels. Importantly, the spatial spread of amplitude modulations increased with time in the low-alpha band but decreased with time in the high-alpha band. Concurrent modulations in high-alpha band amplitude therefore cannot explain the observed high-alpha band phase synchronization.

### Low-alpha wPLI suppression is correlated with low-alpha amplitude suppression across participants and cortical parcels

The low-alpha band desynchronization observed with wPLI was concurrent to the low-alpha amplitude suppression. We investigated whether this decrease in low-alpha band synchronization could be explained by a decrease in SNR due to a concurrent decrease in low-alpha amplitudes. If this were the case, in a given parcel, a stronger decrease in oscillation amplitudes should be associated with a stronger decrease in parcel synchronization strength *PS* (Inline Supplementary Fig. 8). The average correlation coefficients between *PS* and parcel oscillation amplitude were significantly different from 0 (*N* = 13, one-sample t-test, α = 0.05, Holms-Bonferroni corrected; 400–600 ms: *t*(12) = 5.4, *p_corr_
* = 0.00032; 600-800 ms: *t*(12) = 5.0, *p_corr_
* = 0.00032; 800-1000 ms: *t*(12) = 5.0, *p_corr_
* = 0.000086). Across cortical parcels, a larger decrease in oscillation amplitudes was associated with a larger decrease in wPLI values. The suppression of low-alpha band phase-synchronization observed in wPLI could thus not reflect true modulations of neuronal synchronization but rather be explained by SNR suppression.

### High alpha phase synchronization and low-alpha amplitudes display different patterns of hemispheric lateralization

To understand whether high-alpha synchrony and low-alpha amplitudes have a similar role in the implementation of visuospatial attention, we investigated whether high-alpha phase synchronization was lateralized similarly to low-alpha local amplitudes. For each participant, time-frequency bin, and attend condition, we computed ipsi- and contralateral intra-hemispheric graph strength (*GS*) of high-alpha band synchronization and average low-alpha baseline-corrected amplitudes in ipsi- and contralateral visual cortex (Inline Supplementary Fig. 3). High-alpha band GS increased in both ipsi- and contralateral hemispheres regardless of the direction of attention (Fig. 5A). In the attend right condition, GS increased in both ipsi- and contralateral hemispheres concurrently from around 400 ms after cue onset, but the increase was stronger in the ipsilateral (right) hemisphere compared to the contralateral (left) hemisphere. In the attend-left condition, however, an early increase in ipsilateral (left) hemisphere GS was followed by a later increase in contralateral (right) hemisphere GS. GS thus appeared to be stronger in contra-compared to ipsilateral cortex as it started decreasing earlier in the ipsilateral hemisphere. In summary, the lateralization patterns of high-alpha band phase synchronization differed between attend conditions and TWs. In contrast, the pattern of low-alpha local amplitude suppression in ipsi- and contralateral visual cortex was similar for both attend conditions (Fig. 5B). After an initial event-related increase, low-alpha amplitudes decreased in both ipsi- and contralateral visual cortices with this suppression being stronger in contra-compared to ipsilateral visual cortex.

**Figure 5.**
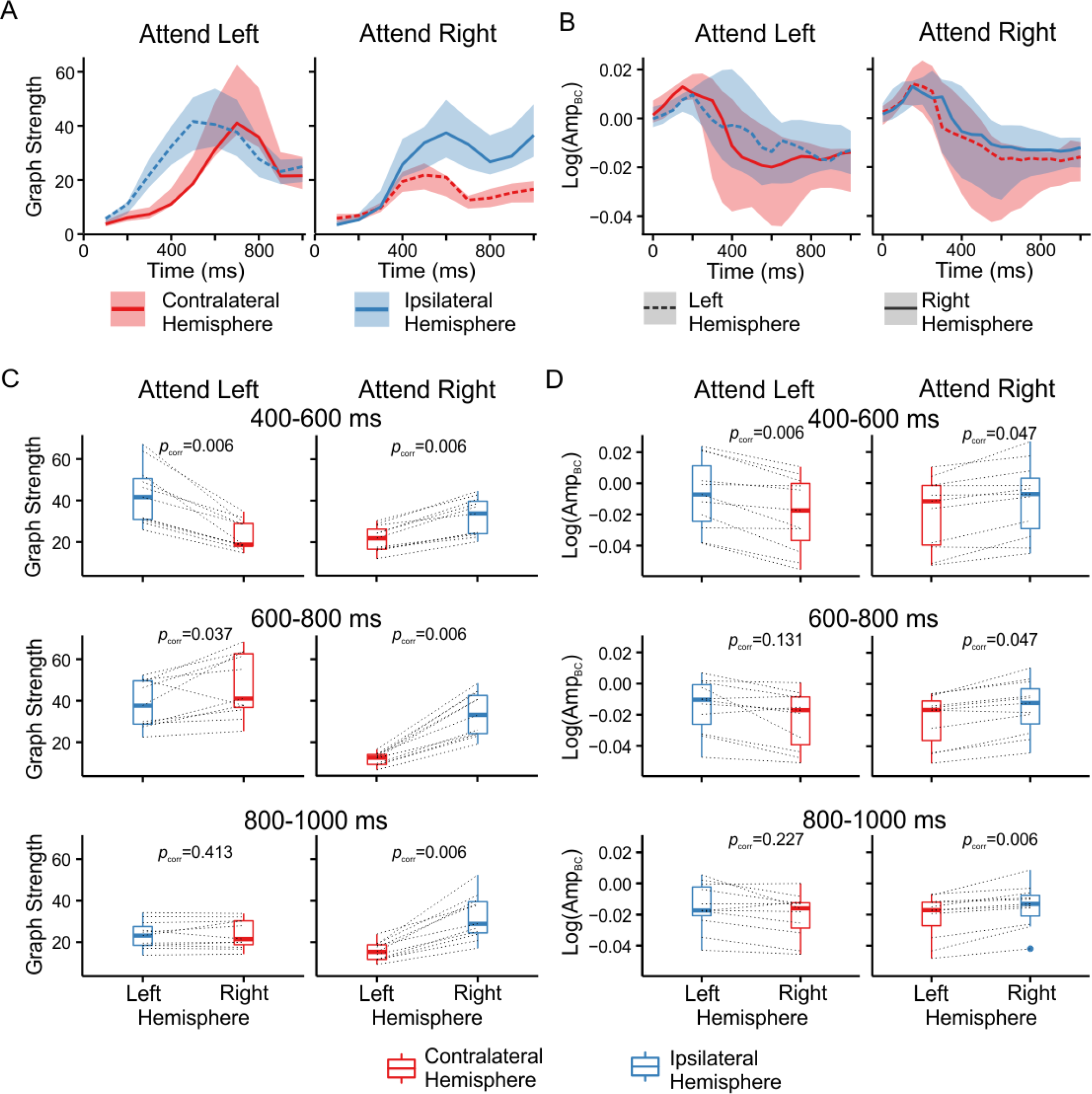
*Lateralization patterns differ between high-alpha inter-areal synchrony and low-alpha local amplitudes*. A: Mean high-alpha graph strength across participants in ipsi- and contralateral hemispheres for the attend-right and attend-left conditions. Shaded areas represent 95% confidence intervals. B: Mean low-alpha amplitudes across participants in visual cortex lateralization in ipsi- and contralateral visual cortices for the attend-right and attend-left conditions. Shaded areas represent 95% confidence intervals. C: Box plots of high-alpha graph strength across participants in the left or right hemisphere for each time-window and attend condition. Dotted lines link individual participant’s data points. Exact multiple-comparison corrected *p* values are given for each comparison. D: Box plots of low-alpha amplitude suppression across participants in the left or right visual cortex for each time-window and attend condition. Dotted lines link individual participant’s data points. Exact multiple-comparison corrected *p* values are given for each comparison.

To further investigate these lateralization phenomena, we tested whether high-alpha *GS* or low-alpha band amplitudes averaged across visual cortex differed between ipsi- and contralateral hemispheres. We ran separate non-parametric tests for each time-window and attend condition (*N* = 11, sign-tests for amplitude data and Wilcoxon signed rank for graph strength, α = 0.05, Holms-Bonferroni corrected). For high-alpha phase synchronization, the pattern of results differed across TWs (Fig. 5C). In the first TW, *GS* was lateralized according to the direction of attention with significantly higher *GS* in the ipsilateral compared to the contralateral hemisphere for both attend conditions. In the second TW, *GS* was significantly stronger in the right hemisphere regardless of the direction of attention. In the third TW, *GS* was significantly stronger in the right hemisphere only in the attend-right condition. As expected, low-alpha amplitude suppression was stronger in contralateral compared to ipsilateral visual cortex for all TWs in the attend-right condition. In the attend-left condition, stronger low-alpha local amplitude suppression in contra-compared to ipsilateral visual cortex was present only in the first TW while there was no difference in suppression strength in last two TWs. While low alpha-band amplitude modulations were consistently stronger in the ipsi- than contralateral in visual cortex, high-alpha band large scale synchronization was clearly lateralized according to the attended hemifield only during the first time-window.

To identify putative differences in the lateralization of phase synchrony within different parts of the networks, we investigated the lateralization of connectivity within the visual system and between the visual system and DAN. We thus counted the number of significant connections for high-alpha band synchronization networks separately within ipsi- and contralateral visual cortex as well as between ipsi- and contralateral visual cortex and bilateral DAN for each attend-condition and time-window. We then computed a lateralization index LI as 
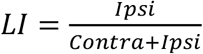
, where values above or below 0.5 indicate respectively more or less connectivity in the ipsilateral compared to the contralateral hemifield. These values were then compared to null-hypothesis values obtained by random shuffling of the networks. All values as well as 95% confidence limits of the null-hypothesis are displayed in Fig. 6. Connectivity in the visual system displayed a similar lateralized pattern in both attend conditions with more connectivity in ipsi-compared to contralateral visual cortex. In contrast, connectivity between the visual system and DAN did not display any such a consistent lateralized pattern across attend-conditions. In summary, while high-alpha band large-scale synchronization was lateralized according to the direction of attention within the visual system, this was not the case for connectivity between the visual system and DAN.

**Figure 6.**
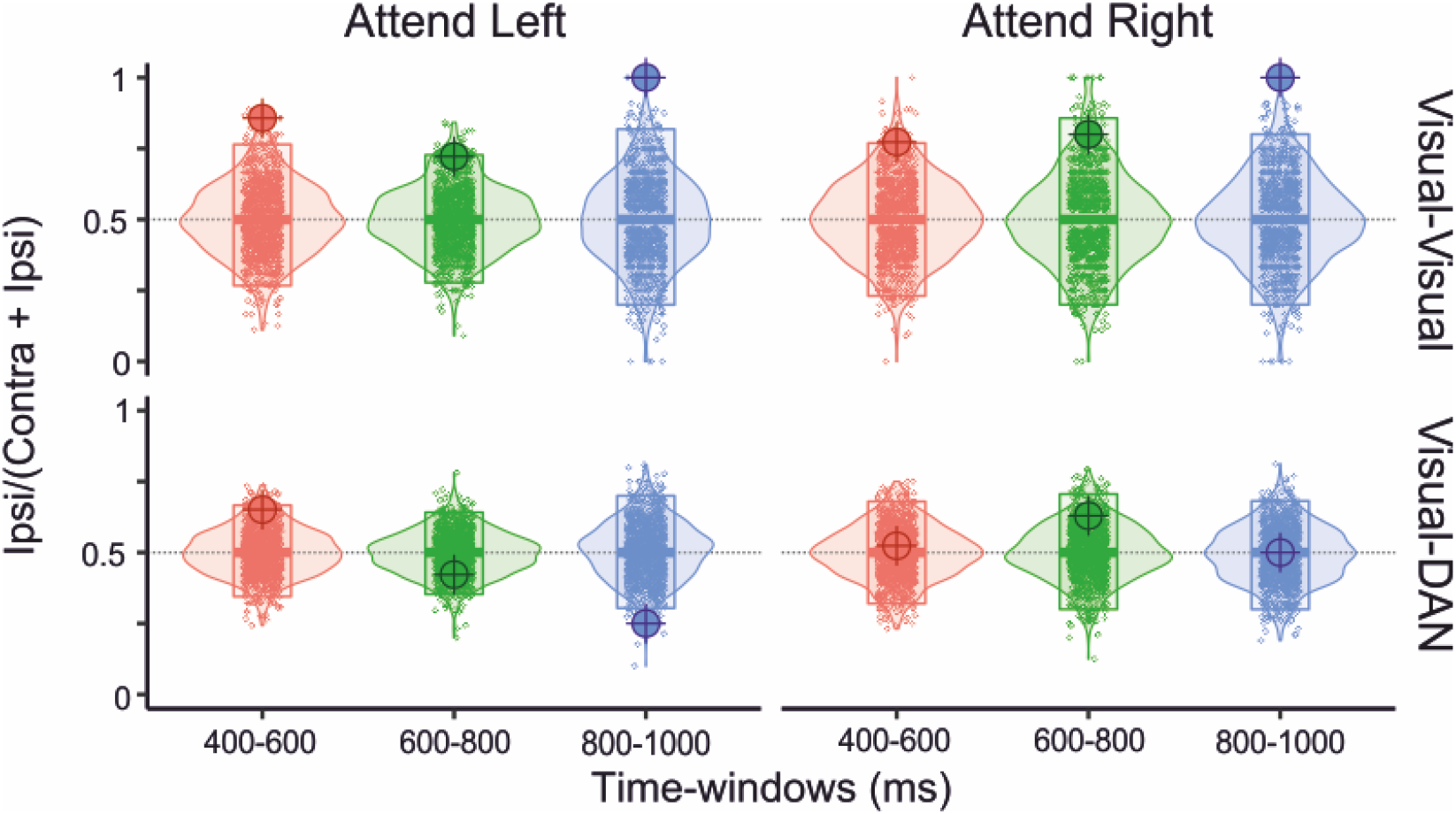
*Lateralization of the number of connections in high-alpha phase synchronization networks*. The lateralization index Ipsi/(Contra+Ipsi) of the number of connections within visual cortex (top row) or between visual cortex and DAN (bottom row) is plotted as a function of attend condition and time-window. For each condition, the null-hypothesis values for 1000 shuffled matrices are displayed over its smoothed distribution shape and a crossbar representing its .025 and .975 quantiles. Lateralization of the number of connections is deemed to be significantly different from that expected by chance if it falls outside this 95% confidence limits.

### Attentional modulations of high-alpha band inter-areal phase synchrony and visual cortex low-alpha oscillation amplitudes co-vary

High-alpha band synchronization should co-vary with the interhemispheric balance of local low-alpha band suppression if it were to mediate attentional effects. We therefore tested whether individual GS co-varied with the lateralization of local low-alpha amplitudes in visual cortex (Fig. 7). We computed an amplitude lateralization index (*ALI*) for each participant as the difference in average baseline-corrected amplitudes between contra- and ipsilateral visual cortex. A more negative ALI indicates stronger alpha suppression in contralateral visual cortex. We then collapsed data for the attend-right and attend-left conditions in each TW and modeled the relationship between GS and ALI using linear mixed models in order to account for the presence of non-independent observations. For the first TW, stronger *GS* was associated with stronger lateralization of amplitude suppression (*N* = 13, *χ^2^
*(1) = 10.2, *p_corr_
* = 0.004, *R^2^
_m_
* = 0.35). For the second TW, *GS* was significantly associated with stronger lateralization of amplitude suppression before correction for multiple comparisons (*N* = 13, *χ^2^
*(1) = 4.3, *p* = 0.039, *R^2^
_m_
* = 0.16, (*p_corr_
* = 0.077). In the last TW, there was no significant relationship between *GS* and amplitude suppression lateralization (*N* = 13, *χ^2^
*(1) = 3.2, *p* = 0.075, *R^2^
_m_
* = 0.11).

**Figure 7.**
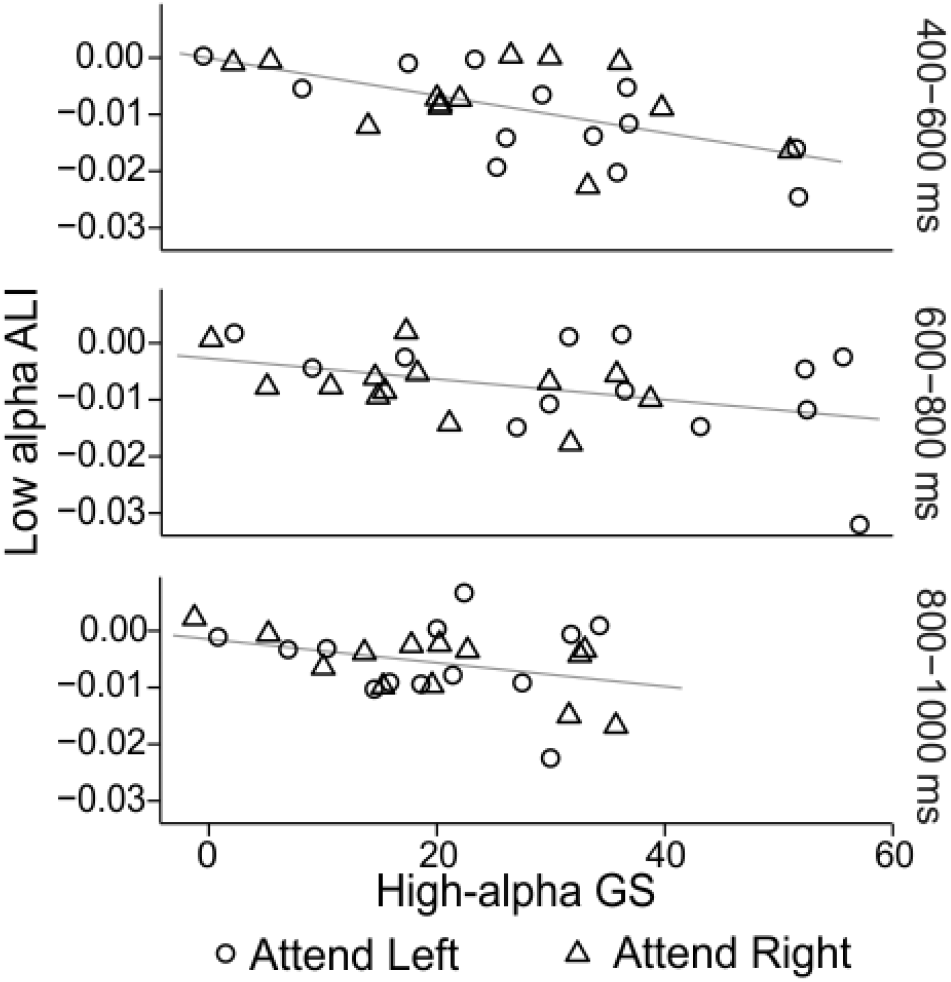
*Network strength co-varies with the lateralization of low-alpha band amplitudes*. Graph strength (GS) of high-alpha band synchronization plotted as a function of the low alpha-band amplitude lateralization index (ALI) across participants for each attend-condition and time-window. Grey lines represent the best fit linear regression line for the fixed effect of GS.

### Strength of phase synchronization predicts behavioral performance

To investigate whether high-alpha band phase-synchronization had an effect on task performance, we tested whether stronger high-alpha band phase synchronization, averaged over the three time-windows of interest, was associated with a stronger benefit of visuospatial attention on RTs. In the low contrast condition (Fig. 8A), there was a strong and significant positive correlation between average *GS* and *RT_benefit_
* for the attend-right condition (*N* = 13, *r* = 0.79, 98.75% *CI* = [0.25 0.96] *p_corr_
* = 0.005; *ρ* = 0.74, 98.75% *CI* = [-.07 0.99] *p_corr_
* = 0.02) but not for the attend-left condition, (*N* = 13, *r* = 0.20, 98.75% *CI* = [-0.80 0.84] *p_corr_
* = 1; *ρ* = 0.09, 98.75% *CI* = [-.82 0.86] *p_corr_
* = 1). In the high contrast condition (Fig. 8B), correlations were of small (attend-left, *N* = 13, *r* = 0.29, 98.75% *CI* = [-0.61 0.81] *p_corr_
* = 1; *ρ* = 0.06, 98.75% *CI* = [-.75 0.80] *p_corr_
* = 1).) or medium (attend-right, *N* = 13, *r* = 0.41, 98.75% *CI* = [-0.55 0.87] *p_corr_
* = 1; *ρ* = 0.29, 98.75% *CI* = [-.74 0.86]*p_corr_
* = 1)) strength but non-significant. In the attend-right condition for low contrast stimuli, participants with stronger average *GS* thus had larger RT differences induced by visuospatial attention than subjects with weaker average *GS*, which shows that inter-areal high-alpha band synchronization was functionally significant.

**Figure 8.**
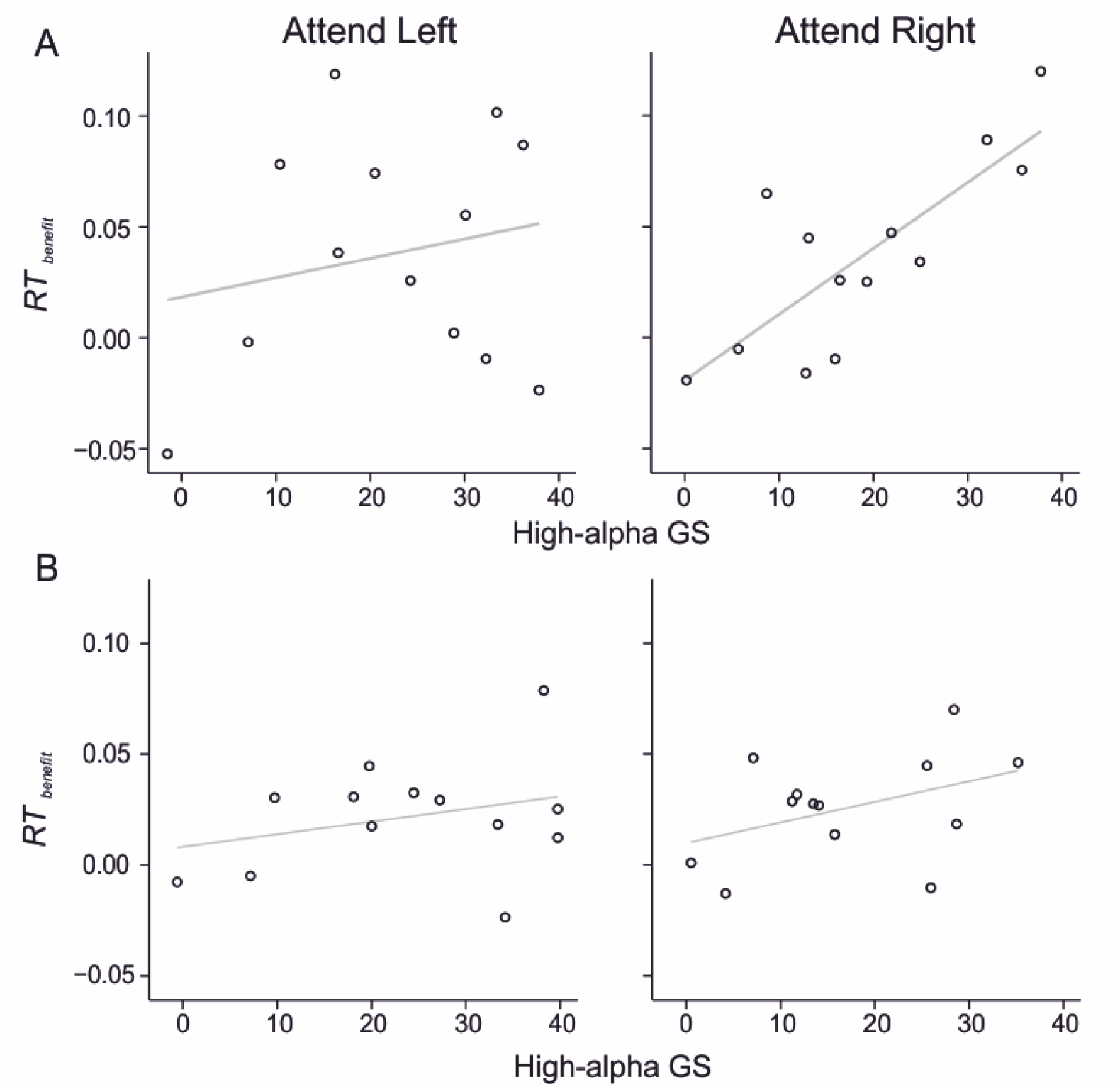
*Relationship between Inter-areal alpha synchronization strength and the behavioral benefit of visuospatial attention*. Scatterplots display *RT_benefit_
* as a function of high-alpha Graph Strength across participants and attention conditions for Low (top row) and High (bottom row) contrasts. RT_benefit_ was estimated as the standardized difference between RTs for valid and invalid trials. Grey lines indicate the best fit linear regression line.

## Discussion

We used data-driven analyses of source-localized MEG data to assess the role of large-scale phase synchronization of cortical oscillations in anticipatory visuospatial attention. Our study reproduced the commonly observed lateralized suppression of local low-alpha amplitudes in visual cortex. This well-known phenomenon was, however, paralleled by a previously unreported and robust strengthening of inter-areal phase synchronization exclusively in the high-alpha frequency band. High-alpha synchronization connected frontoparietal attention systems both to each other and to the visual system. High-alpha band connectivity within the visual system was lateralized according to the direction of attention, similarly to low-alpha amplitude suppression. In contrast, high-alpha band synchronization between visual and attentional systems exhibited lateralization patterns and cortical organization distinct from those of low-alpha amplitude suppression. Crucially, the strength of high-alpha phase-synchronization was correlated with both low-alpha amplitude suppression and attentional task performance benefits, *i.e*., with improved reaction times. These data thus demonstrate that anticipatory visuospatial attention is associated with robust large-scale synchronization in the high-alpha but not in theta or beta/gamma bands and that this synchronization connects the cortical regions previously associated with attentional functions. High-alpha band synchronization could thus play a role in coordinating and regulating neuronal processing across frontoparietal and visual systems and thereby also implement attentional modulations of low-alpha amplitudes.

### Visuospatial attention modulates behavioral performance and low-alpha band amplitudes

Visuospatial attention improved both RTs and detection rates for low-contrast stimuli, consistent with previous research (Capilla et al., 2014; Posner, 1980). For high-visibility supra-threshold stimuli, attention improved RTs but not discrimination performance. Visuospatial attention was further associated with the frequently reported sustained suppression of local alpha- and beta-band amplitudes (Capilla et al., 2014; Liu et al., 2016; Thut et al., 2006). In line with prior results when both attended and non-attended stimuli were to be reported, this amplitude suppression, albeit stronger in the contralateral visual cortex, was present in both ipsi- and contralateral visual cortices (Gould et al., 2011). These results thus validate that participants were covertly attending to the cued hemifield and reproduce prior observations of lateralized alpha amplitude suppression.

### Inter-areal high-alpha band phase synchronization characterizes visuospatial attention

Anticipatory visuospatial attention was associated with a robust and sustained strengthening of inter-areal phase synchronization in the high-alpha frequency band only. Importantly, visuospatial attention did not modulate phase synchrony in any of the other studied frequency bands. Several characteristics of strengthened high-alpha synchrony were consistent with its putative functional role in visuospatial attention. This strengthening started shortly after cue-onset, once cue information had been extracted (Simpson et al., 2011) but preceded the low-alpha amplitude suppression by 100-200 ms. High-alpha synchronization might thus first support the shifting of attention to the cued hemifield (first TW). It was also sustained throughout the post-cue interval, suggesting that it could later (second and third TW) support the maintenance of attention to the cued hemifield (Simpson et al., 2011). Furthermore, similarly to local amplitudes, lateralization of the synchrony networks in the visual system was dependent on the direction of attention, indicating that strengthened high-alpha synchrony was related to visuospatial attention task demands. Importantly, there was no concurrent increase in high-alpha amplitudes or stimulus locking, which indicates that strengthened synchronization was not attributable to changes in the signal-to-noise ratio or to cue-onset-locked phenomena such as alpha-ringing (Hindriks et al., 2014; Makeig et al., 2002). These results thus yield strong evidence for the overarching hypothesis that alpha-band long-range phase synchronization regulates inter-areal communication to coordinate attentional modulations (Palva and Palva, 2007; Palva and Palva, 2011; Sadaghiani and Kleinschmidt, 2016).

Our results expand prior studies showing that anticipatory attention (Doesburg et al., 2009; Sacchet et al., 2015; Siegel et al., 2008) is associated with phase synchronization in alpha and beta / gamma bands in EEG/MEG data and in delta (1-4 Hz) and theta bands in human intracranial EEG data (Daitch et al., 2013). Importantly, in contrast to prior human non-invasive studies (Siegel et al., 2008) and non-human primate invasive LFP studies (Gregoriou et al., 2009; Womelsdorf et al., 2007), we did not observe attention-dependent synchronization in the beta or gamma bands. These inconsistencies could result from differences in experimental paradigms. Primate studies investigated attention in the presence of to-be-attended visual stimuli whereas in our paradigm there was no stimulus displayed during the attentional delay. Furthermore, while prior MEG and EEG studies only investigated the attention-dependent lateralization of phase synchronization, our study addressed the complete cortical attention phase synchronized-networks.

Network synchronization was not fully lateralized according to the direction of attention and differed between the switching and maintenance of visual attention. As in prior studies of visuospatial attention (Doesburg et al., 2009; Siegel et al., 2008), high-alpha band synchronization was stronger in the ipsilateral hemisphere during attentional shifting, in line with interhemispheric balance models of visual attention (Buschman and Kastner, 2015; Scolari et al., 2015; Szczepanski et al., 2013). However, during the subsequent attentional maintenance, high-alpha band networks became right lateralized regardless of attended hemifield, which supports models of right-hemispheric dominance of attention (Heilman and Van Den Abell, 1980; Zago et al., 2016). Right-hemispheric lateralization could be driven by the presence of phase synchrony between right frontal cortex and DAN or sensory systems as previously reported (Sacchet et al., 2015) and favored by right-lateralized structural connectivity (Marshall et al., 2015a). Importantly, the lateralization of network synchronization differed across systems. In the visual system, there was more connectivity in the ipsi- compared to the contralateral hemifield. Synchrony was thus lateralized according to the direction of attention similar to local alpha amplitudes (Gould et al., 2011; Liu et al., 2016; Thut et al., 2006). In contrast, connectivity between the visual system and DAN did not display any consistent lateralized pattern and, when considering the networks with all cortical systems, high-alpha band synchronization was right-lateralized. Alpha-band synchronization within the visual system and between the visual system and attention networks therefore reflect distinct processes in the implementation of visuospatial attention.

### Dynamic high-alpha band synchronization connects nodes of attentional and visual systems

Key hubs of the networks of high-alpha phase synchronization were observed in cortical systems previously shown to be involved in attentional processing. Central hubs were observed in key nodes of DAN and FPN (IPS, SPG, right IFG and MFS) that exhibit increased BOLD activity (Corbetta and Shulman, 2002; Kastner and Ungerleider, 2000) and peaks in event-related fields (Simpson et al., 2011) during anticipatory visual attention. Network hubs were also found across the visual processing hierarchy from V1 to the dorsal and ventral stream areas that represent visual object information (Kravitz et al., 2013), as expected if visuospatial attention modulates cortical activity at several levels of the visual processing hierarchy (Buffalo et al., 2010; Capilla et al., 2014).

High-alpha phase synchrony connected cortical areas not only within DAN and visual systems but also between DAN and FPN as well as between DAN, FPN and the visual system. The presence of extensive connectivity between these different cortical systems extends previous observations on the role of long-range synchronization in visuospatial attention. Previous studies reported increased connectivity between a few a priori-chosen nodes belonging to FPN, DAN, or the visual system in non-human primate (Buschman and Miller, 2007; Gregoriou et al., 2009), or human EEG/MEG studies (Doesburg et al., 2009; Siegel et al., 2008) but had not investigated the complete cortical networks encompassing these systems. As our analyses did not make assumptions about the possible localization of cortical hubs or inter-areal connectivity, they reveal the most robust connections among all possible connections (Kriegeskorte et al., 2009). Despite this absence of *a priori* choices, the anatomical structure of the observed networks, in terms of both hubs and edges, is consistent with the literature. Furthermore, network anatomical structures were different for attend-right and attend-left conditions, indicating that they cannot solely reflect task demands such as response anticipation or cue processing that remain unchanged between these conditions. Dynamic coupling of visual, FPN, and DAN systems indicates that high-alpha band phase synchronization could play a functional role in visuospatial attention, possibly by facilitating top-down communication (Bastos et al., 2015; Fries, 2015).

### High-alpha band phase synchronization plays a functional role in attention-imposed modulations of behavioral and neuronal activity

High-alpha band phase synchronization co-varied with neuronal and behavioral measures of attention, further supporting the idea of functional relevance to visuospatial attention. Participants who exhibited stronger high-alpha phase synchronization also exhibited stronger lateralization of low-alpha amplitudes during attention shifting. High-alpha phase synchronization between frontal and parietal cortices could thus mediate attention-related low-alpha band suppression, similar to structural connectivity (Marshall et al., 2015a) and perturbation of frontoparietal cortex with TMS (Capotosto et al., 2009; Capotosto et al., 2016; Capotosto et al., 2012; Marshall et al., 2015b). In addition, participants with stronger high-alpha phase synchronization exhibited a stronger behavioral benefit of visuospatial attention when detecting low-contrast but not high contrast stimuli in the right hemifield. This is in line with the hypothesis that phase-synchrony between frontoparietal networks and the visual system could facilitate conscious access to perceptual threshold stimuli (Dehaene and Changeux, 2011) while such synchronization would not modulate discrimination performance for supra-threshold stimuli, which is based on distinct systems-level mechanisms (Sadaghiani and Kleinschmidt, 2016). The correlations of individual GS values with low-alpha amplitude suppression and RT benefit suggest that the identified phase-synchrony networks may be discernible also at a single-subject level. Identifying how individual variability in attention network structures affects attentional processing will be a critical step to understanding how abnormalities in attention networks may lead to attentional deficits in neurodevelopmental disorders or neurological diseases.

Taken together, our results demonstrate that high-alpha band synchronization between frontoparietal and visual systems may underlie the implementation and maintenance of visuospatial attention in humans (Fries, 2015; Womelsdorf and Everling, 2015). More specifically, high-alpha band phase synchronization between multiple key nodes of FPN and DAN could implement the coordination of multiple priority maps distributed across frontal, parietal and collicular structures (Womelsdorf and Everling, 2015).

### Local amplitude dynamics and inter-areal phase coupling play distinct roles in visuospatial attention

The functional role of alpha-band oscillations and inter-areal phase synchrony in cortical computations and attention is debated and different models have been proposed to account for empirical data (Jensen and Mazaheri, 2010; Klimesch et al., 2007; Palva and Palva, 2007; Palva and Palva, 2011; Sadaghiani and Kleinschmidt, 2016). Our results support models where local alpha amplitudes and inter-areal synchronization have distinct functional roles in perception and attention (Bonnefond et al., 2017; Palva and Palva, 2007; Palva and Palva, 2011; Sadaghiani and Kleinschmidt, 2016). The strength of low-alpha amplitudes in visual cortex decreased during visuospatial attention, consolidating the widespread view that suppression of local alpha oscillations in sensory cortices facilitates processing of attended sensory stimuli (Jensen and Mazaheri, 2010; Klimesch et al., 2007; Lange et al., 2013). Similarly, high-alpha connectivity was lateralized according to the direction of attention in the visual cortex, inline with the hypothesis that local alpha-band synchronization is correlated with inhibitory effects. In contrast, the overall strength of high-alpha band phase synchronization increased during the coordination and maintenance of visuospatial attention and did not follow the lateralization pattern classically associated with local alpha amplitudes, arguing against a similar ‘inhibitory’ role for inter-areal alpha phase synchrony between frontoparietal and visual areas (Palva and Palva, 2007; Palva and Palva, 2011). Our results are consistent with a framework where high-alpha synchrony underlies anticipatory endogenous control of sustained visuospatial attention by facilitating communication between relevant cortical areas (Palva and Palva, 2007; Palva and Palva, 2011; Sadaghiani and Kleinschmidt, 2016).

### Conclusion

We found high-alpha phase synchronization to be associated with anticipatory visuospatial attention. High-alpha synchronization connected the major hubs of DAN, FPN and visual systems while synchronization strength predicted both the behavioral attentional benefit and low-alpha amplitude suppression. High-alpha band synchronization could thus support anticipatory endogenous attention by regulating collective neuronal processing across frontal, parietal and visual cortices and modulating local alpha-band amplitudes in the visual cortex.

## Notes

This study was supported by the Academy of Finland (SA 266402 and SA 273807 to S.P. and SA 253130 to J.M.P.). The authors declare no competing financial interests

## References

Bastos, A.M., Vezoli, J., Bosman, C.A., Schoffelen, J.M., Oostenveld, R., Dowdall, J.R., De Weerd, P., Kennedy, H., Fries, P., 2015. Visual areas exert feedforward and feedback influences through distinct frequency channels. Neuron. 85, 390–401.

Bates, D., Mᅡñchler, M., Bolker, B., Walker, S., 2015. Fitting linear mixed-effects models using lme4. Journal of Statistical Software; Vol 1, Issue 1 (2015).

Bonacich, P., 2007. Some unique properties of eigenvector centrality. Social Networks. 29, 555–564.

Bonnefond, M., Kastner, S., Jensen, O., 2017. Communication between brain areas based on nested oscillations. eNeuro. 4.

Brainard, D.H., 1997. The psychophysics toolbox. Spat. Vis. 10, 433–436.

Buffalo, E.A., Fries, P., Landman, R., Liang, H., Desimone, R., 2010. A backward progression of attentional effects in the ventral stream. Proc. Natl. Acad. Sci. U. S. A. 107, 361–365.

Bullmore, E., Sporns, O., 2009. Complex brain networks: Graph theoretical analysis of structural and functional systems. Nat. Rev. Neurosci. 10, 186–198.

Buschman, T.J., Miller, E.K., 2007. Top-down versus bottom-up control of attention in the prefrontal and posterior parietal cortices. Science. 315, 1860–1862.

Buschman, T., Kastner, S., 2015. From behavior to neural dynamics: An integrated theory of attention. Neuron. 88, 127–144.

Capilla, A., Schoffelen, J.M., Paterson, G., Thut, G., Gross, J., 2014. Dissociated alpha-band modulations in the dorsal and ventral visual pathways in visuospatial attention and perception. Cereb. Cortex. 24, 550–561.

Capotosto, P., Babiloni, C., Romani, G.L., Corbetta, M., 2009. Frontoparietal cortex controls spatial attention through modulation of anticipatory alpha rhythms. J. Neurosci. 29, 5863–5872.

Capotosto, P., Baldassarre, A., Sestieri, C., Spadone, S., Romani, G.L., Corbetta, M., 2016. Task and regions specific top-down modulation of alpha rhythms in parietal cortex. Cereb. Cortex.

Capotosto, P., Babiloni, C., Romani, G.L., Corbetta, M., 2012. Differential contribution of right and left parietal cortex to the control of spatial attention: A simultaneous EEG–rTMS study. Cerebral Cortex. 22, 446–454.

Capotosto, P., Spadone, S., Tosoni, A., Sestieri, C., Romani, G.L., Della Penna, S., Corbetta, M., 2015. Dynamics of EEG rhythms support distinct visual selection mechanisms in parietal cortex: A simultaneous transcranial magnetic stimulation and EEG study. The Journal of Neuroscience. 35, 721–730.

Corbetta, M., Shulman, G.L., 2002. Control of goal-directed and stimulus-driven attention in the brain. Nat. Rev. Neurosci. 3, 201–215.

Daitch, A.L., Sharma, M., Roland, J.L., Astafiev, S.V., Bundy, D.T., Gaona, C.M., Snyder, A.Z., Shulman, G.L., Leuthardt, E.C., Corbetta, M., 2013. Frequency-specific mechanism links human brain networks for spatial attention. Proc. Natl. Acad. Sci. U. S. A. 110, 19585–19590.

Dale, A.M., Liu, A.K., Fischl, B.R., Buckner, R.L., Belliveau, J.W., Lewine, J.D., Halgren, E., 2000. Dynamic statistical parametric mapping: Combining fMRI and MEG for high-resolution imaging of cortical activity. Neuron. 26, 55–67.

Dehaene, S., Changeux, J.P., 2011. Experimental and theoretical approaches to conscious processing. Neuron. 70, 200–227.

Destrieux, C., Fischl, B., Dale, A., Halgren, E., 2010. Automatic parcellation of human cortical gyri and sulci using standard anatomical nomenclature. Neuroimage. 53, 1–15.

Doesburg, S.M., Green, J.J., McDonald, J.J., Ward, L.M., 2009. From local inhibition to long-range integration: A functional dissociation of alpha-band synchronization across cortical scales in visuospatial attention. Brain Res. 1303, 97–110.

Fischl, B., 2012. FreeSurfer. Neuroimage. 62, 774–781.

Fries, P., 2015. Rhythms for cognition: Communication through coherence. Neuron. 88, 220–235.

Gould, I.C., Rushworth, M.F., Nobre, A.C., 2011. Indexing the graded allocation of visuospatial attention using anticipatory alpha oscillations. J. Neurophysiol. 105, 1318–1326.

Gramfort, A., Luessi, M., Larson, E., Engemann, D.A., Strohmeier, D., Brodbeck, C., Parkkonen, L., Hämäläinen, M.S., 2014. MNE software for processing MEG and EEG data. Neuroimage. 86, 446–460.

Gregoriou, G.G., Gotts, S.J., Zhou, H., Desimone, R., 2009. High-frequency, long-range coupling between prefrontal and visual cortex during attention. Science. 324, 1207–1210.

Heilman, K.M., Van Den Abell, T., 1980. Right hemisphere dominance for attention: The mechanism underlying hemispheric asymmetries of inattention (neglect). Neurology. 30, 327–330.

Hindriks, R., van Putten, M.J., Deco, G., 2014. Intra-cortical propagation of EEG alpha oscillations. Neuroimage. 103, 444–453.

Iemi, L., Chaumon, M., Crouzet, S.M., Busch, N.A., 2017. Spontaneous neural oscillations bias perception by modulating baseline excitability. J. Neurosci. 37, 807–819.

Jensen, O., Mazaheri, A., 2010. Shaping functional architecture by oscillatory alpha activity: Gating by inhibition. Front. Hum. Neurosci. 4, 186.

Kastner, S., Ungerleider, L.G., 2000. Mechanisms of visual attention in the human cortex. Annu. Rev. Neurosci. 23, 315–341.

Klimesch, W., Sauseng, P., Hanslmayr, S., 2007. EEG alpha oscillations: The inhibition-timing hypothesis. Brain Res. Rev. 53, 63–88.

Korhonen, O., Palva, S., Palva, J.M., 2014. Sparse weightings for collapsing inverse solutions to cortical parcellations optimize M/EEG source reconstruction accuracy. J. Neurosci. Methods. 226C, 147–160.

Kravitz, D.J., Saleem, K.S., Baker, C.I., Ungerleider, L.G., Mishkin, M., 2013. The ventral visual pathway: An expanded neural framework for the processing of object quality. Trends Cogn. Sci. 17, 26–49.

Kriegeskorte, N., Simmons, W.K., Bellgowan, P.S., Baker, C.I., 2009. Circular analysis in systems neuroscience: The dangers of double dipping. Nat. Neurosci. 12, 535–540.

Lachaux, J.P., Rodriguez, E., Martinerie, J., Varela, F.J., 1999. Measuring phase synchrony in brain signals. Hum. Brain Mapp. 8, 194–208.

Lange, J., Oostenveld, R., Fries, P., 2013. Reduced occipital alpha power indexes enhanced excitability rather than improved visual perception. J. Neurosci. 33, 3212–3220.

Liu, Y., Bengson, J., Huang, H., Mangun, G.R., Ding, M., 2016. Top-down modulation of neural activity in anticipatory visual attention: Control mechanisms revealed by simultaneous EEG-fMRI. Cerebral Cortex. 26, 517–529.

Makeig, S., Westerfield, M., Jung, T.P., Enghoff, S., Townsend, J., Courchesne, E., Sejnowski, T.J., 2002. Dynamic brain sources of visual evoked responses. Science. 295, 690–694.

Marshall, T.R., Bergmann, T.O., Jensen, O., 2015a. Frontoparietal structural connectivity mediates the top-down control of neuronal synchronization associated with selective attention. PLoS Biol. 13, e1002272.

Marshall, T.R., O’Shea, J., Jensen, O., Bergmann, T.O., 2015b. Frontal eye fields control attentional modulation of alpha and gamma oscillations in contralateral occipitoparietal cortex. The Journal of Neuroscience. 35, 1638–1647.

Miller, E.K., Buschman, T.J., 2013. Cortical circuits for the control of attention. Curr. Opin. Neurobiol. 23, 216–222.

Oostenveld, R., Fries, P., Maris, E., Schoffelen, J.M., 2011. FieldTrip: Open source software for advanced analysis of MEG, EEG, and invasive electrophysiological data. Comput. Intell. Neurosci. 2011, 156869.

Palva, J.M., Monto, S., Kulashekhar, S., Palva, S., 2010. Neuronal synchrony reveals working memory networks and predicts individual memory capacity. Proc. Natl. Acad. Sci. U. S. A. 107, 7580–7585.

Palva, S., Palva, J.M., 2007. New vistas for alpha-frequency band oscillations. Trends Neurosci. 30, 150–158.

Palva, S., Palva, J.M., 2011. Functional roles of alpha-band phase synchronization in local and large-scale cortical networks. Front. Psychol. 2, 204.

Palva, S., Palva, J.M., 2012. Discovering oscillatory interaction networks with M/EEG: Challenges and breakthroughs. Trends Cogn. Sci. 16, 219–230.

Palva, S., Kulashekhar, S., Hamalainen, M., Palva, J.M., 2011. Localization of cortical phase and amplitude dynamics during visual working memory encoding and retention. J. Neurosci. 31, 5013–5025.

Pernet, C.R., Wilcox, R.R., Rousselet, G.A., 2013. Robust correlation analyses: False positive and power validation using a new open source matlab toolbox. Frontiers in Psychology. 3, 606.

Petersen, S.E., Posner, M.I., 2012. The attention system of the human brain: 20 years after. Annu. Rev. Neurosci. 35, 73–89.

Posner, M.I., 1980. Orienting of attention. Q J Exp Psychol. 32, 3–25.

Rouhinen, S., Panula, J., Palva, J.M., Palva, S., 2013. Load dependence of beta and gamma oscillations predicts individual capacity of visual attention. J. Neurosci. 33, 19023–19033.

Rubinov, M., Sporns, O., 2010. Complex network measures of brain connectivity: Uses and interpretations. Neuroimage. 52, 1059–1069.

Sacchet, M.D., LaPlante, R.A., Wan, Q., Pritchett, D.L., Lee, A.K.C., Hämäläinen, M., Moore, C.I., Kerr, C.E., Jones, S.R., 2015. Attention drives synchronization of alpha and beta rhythms between right inferior frontal and primary sensory neocortex. The Journal of Neuroscience. 35, 2074–2082.

Sadaghiani, S., Kleinschmidt, A., 2016. Brain networks and α-oscillations: Structural and functional foundations of cognitive control. Trends Cogn. Sci. (Regul. Ed.). 20, 805–817.

Schoffelen, J.M., Gross, J., 2009. Source connectivity analysis with MEG and EEG. Hum. Brain Mapp. 30, 1857–1865.

Scolari, M., Seidl-Rathkopf, K.N., Kastner, S., 2015. Functions of the human frontoparietal attention network: Evidence from neuroimaging. Current Opinion in Behavioral Sciences. 1, 32–39.

Siegel, M., Donner, T.H., Oostenveld, R., Fries, P., Engel, A.K., 2008. Neuronal synchronization along the dorsal visual pathway reflects the focus of spatial attention. Neuron. 60, 709–719.

Simpson, G.V., Weber, D.L., Dale, C.L., Pantazis, D., Bressler, S.L., Leahy, R.M., Luks, T.L., 2011. Dynamic activation of frontal, parietal, and sensory regions underlying anticipatory visual spatial attention. J. Neurosci. 31, 13880–13889.

Spadone, S., Della Penna, S., Sestieri, C., Betti, V., Tosoni, A., Perrucci, M.G., Romani, G.L., Corbetta, M., 2015. Dynamic reorganization of human resting-state networks during visuospatial attention. Proc. Natl. Acad. Sci. U. S. A. 112, 8112–8117.

Szczepanski, S.M., Konen, C.S., Kastner, S., 2010. Mechanisms of spatial attention control in frontal and parietal cortex. The Journal of Neuroscience: The Official Journal of the Society for Neuroscience. 30, 148–160.

Szczepanski, S.M., Pinsk, M.A., Douglas, M.M., Kastner, S., Saalmann, Y.B., 2013. Functional and structural architecture of the human dorsal frontoparietal attention network. Proc. Natl. Acad. Sci. U. S. A. 110, 15806–15811.

Tallon-Baudry, C., 2012. On the neural mechanisms subserving consciousness and attention. Front. Psychol. 2, 397.

Thut, G., Nietzel, A., Brandt, S.A., Pascual-Leone, A., 2006. Alpha-band electroencephalographic activity over occipital cortex indexes visuospatial attention bias and predicts visual target detection. J. Neurosci. 26, 9494–9502.

van Dijk, H., Schoffelen, J.M., Oostenveld, R., Jensen, O., 2008. Prestimulus oscillatory activity in the alpha band predicts visual discrimination ability. J. Neurosci. 28, 1816–1823.

Vinck, M., Oostenveld, R., van Wingerden, M., Battaglia, F., Pennartz, C.M., 2011. An improved index of phase-synchronization for electrophysiological data in the presence of volume-conduction, noise and sample-size bias. Neuroimage. 55, 1548–1565.

Watson, A.B., Pelli, D.G., 1983. QUEST: A bayesian adaptive psychometric method. Percept. Psychophys. 33, 113–120.

Womelsdorf, T., Everling, S., 2015. Long-range attention networks: Circuit motifs underlying endogenously controlled stimulus selection. Trends Neurosci. 38, 682–700.

Womelsdorf, T., Fries, P., 2007. The role of neuronal synchronization in selective attention. Curr. Opin. Neurobiol. 17, 154–160.

Womelsdorf, T., Schoffelen, J.M., Oostenveld, R., Singer, W., Desimone, R., Engel, A.K., Fries, P., 2007. Modulation of neuronal interactions through neuronal synchronization. Science. 316, 1609–1612.

Yeo, B.T., Krienen, F.M., Sepulcre, J., Sabuncu, M.R., Lashkari, D., Hollinshead, M., Roffman, J.L., Smoller, J.W., Zollei, L., Polimeni, J.R., Fischl, B., Liu, H., Buckner, R.L., 2011. The organization of the human cerebral cortex estimated by intrinsic functional connectivity. J. Neurophysiol. 106, 1125–1165.

Zago, L., Petit, L., Jobard, G., Hay, J., Mazoyer, B., Tzourio-Mazoyer, N., Karnath, H.O., Mellet, E., 2016. Pseudoneglect in line bisection judgement is associated with a modulation of right hemispheric spatial attention dominance in right-handers. Neuropsychologia. 94, 75–83.

